# Development of a novel peripherally acting alpha2A-adrenergic receptor antagonist for anti-diabetic

**DOI:** 10.1101/2024.12.17.628818

**Authors:** Feng Hong, Jingyi Wang, Dongsheng Chen, Qian Wang, Shiyi Gan, Fang Ye, Lihua Chen, Huixia Ren, Tongran Zhang, Yinfang Xie, Xiaoyuan Jing, Ruoxi Wang, Zhibin Xu, Jiayan Ren, Cuihan Shi, Shuhui Jia, Xiaoting Huang, Xiawei Xu, Jing Lu, Yunhua Zheng, Wanhua Lin, Ji Dai, Liping Wang, Liangyi Chen, Chao Tang, Huisheng Liu, Tao Xu, Xin Xie, Xin-an Liu, Yang Du, Fajun Nan, Zuxin Chen

## Abstract

Yohimbine, a potent alpha2A-adrenergic receptor (α_2A_AR) antagonist, was found therapeutic potential for type 2 diabetes through improving insulin release. However, the adverse side effects mediated by its actions in the brain hampered its use. Here, based on molecular docking analysis and structural modification, we have developed a novel peripherally acting yohimbine derivative (CDS479-2). CryoEM data found that yohimbine and CDS479-2 have similar interactions with the structure of α_2A_AR. Importantly, CDS479-2 shows similar α_2A_AR antagonist activity as yohimbine, but with very limited access to the brain, and thus avoiding the unwanted central effects such as hypertension and anxiety. Acute administration of CDS479-2 by injection or gavage lowered blood glucose levels and improved glucose tolerance in the high-fat diet-induced obesity (DIO) mice, an animal model for human type 2 diabetes. Remarkably, DIO mice received 2 weeks of daily administration of CDS479-2, but not yohimbine, exhibited sustained normoglycaemia, and increased density of the insulin-producing beta cells, in which important proliferation genes were found upregulated. Moreover, the overall protein expression levels of their pancreas were more similar to that of the healthy chow-fed mice. Thus, CDS479-2 may indicate a new direction for type 2 diabetes treatment. Importantly, the strategy we employed in this study will inspire the optimization for drugs that with both peripheral and central targets.

**Graphic abstract:** 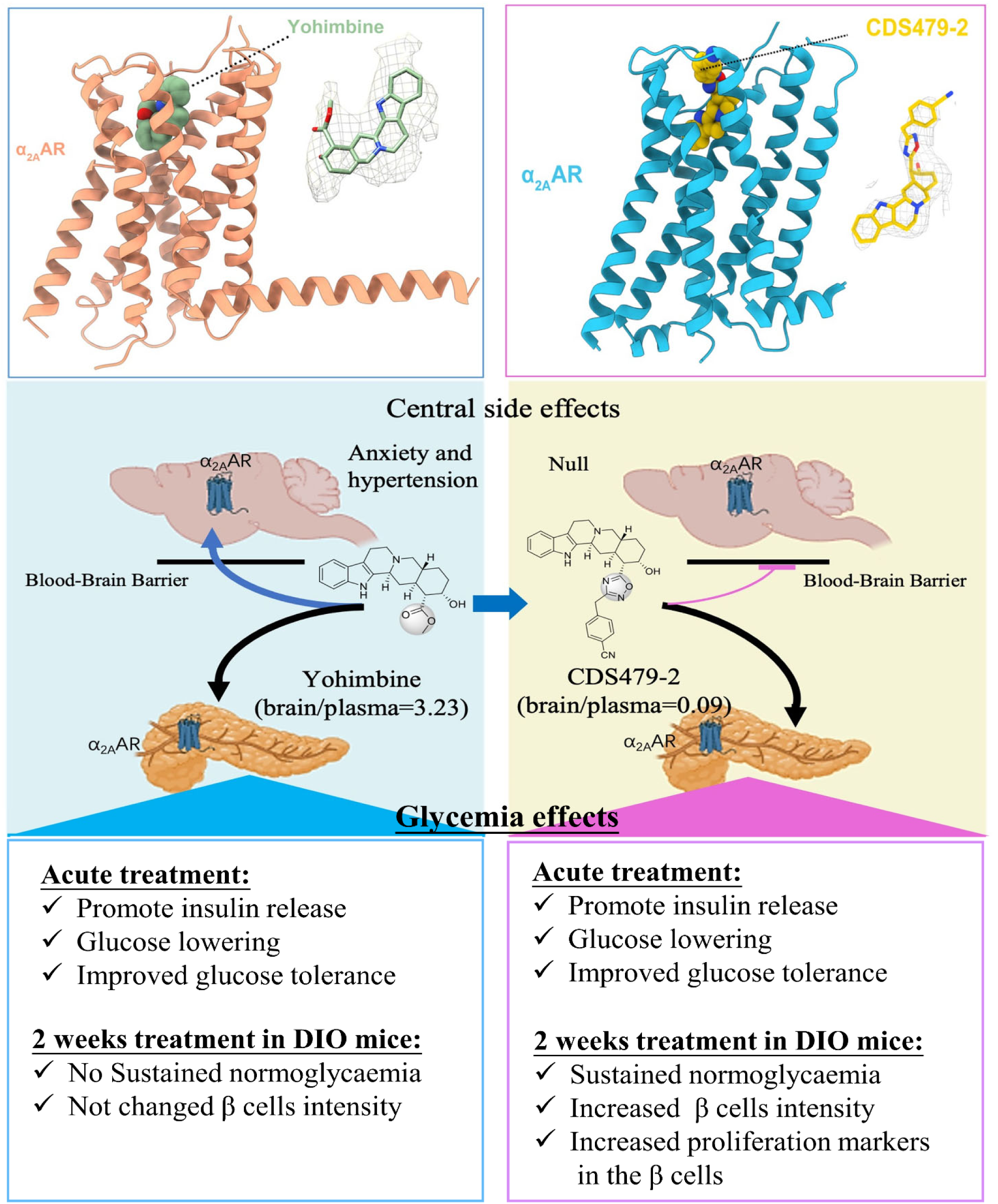

## Introduction

Genetic variants in the gene *ADRA2A*, which encodes for the alpha2A-adrenergic receptor (α_2A_AR), were consistently found associated with lowered plasma insulin levels and increased type 2 diabetes risk ^1–7^. Previous studies have demonstrated that pancreatic islets from the risk variants carriers displayed an overexpression of α_2A_AR, impaired insulin secretion, and dysfunction of insulin-producing β cells^1,7^. α_2A_AR belongs to the GPCR superfamily and commonly couples to the inhibitory GTP–binding proteins (Gi proteins) ^8^. Activation of α_2A_AR in the pancreatic β-cells was found suppressing insulin release and elevating blood glucose levels ^9,10^. Thus, the impairment of insulin secretion observed in the *ADRA2A* risk variant carrier was probably caused by the over-activity of α_2A_AR, and antagonism of α_2A_AR would improve insulin release. Indeed, yohimbine, a commonly used α_2A_AR antagonist in clinical and pre-clinical studies, was found to correct the defects of insulin secretion and elevate insulin levels in the type 2 diabetes patients who carrying the risk variant^1^. Thus, yohimbine was suggested as a therapeutic potential mean for these patients, which accounts for 40% of type 2 diabetes patients of the Caucasian origin^7^. However, given α_2A_AR widely distributed in the central nervous system, and *ADRA2A* polymorphisms were also associated with hypertension and depression-related suicide, manipulation of α_2A_AR signaling would probably affect the brain functions ^8,11,12^. Indeed, acute yohimbine treatment was found elevated systolic blood pressures and anxiety levels in the clinical trial^1^. These adverse side effects were likely mediated by the actions of yohimbine on the central α_2A_AR located in the brain areas that regulate mood and autonomic functions^12–16^. Thus, development of peripherally acting α_2A_AR antagonists to improve insulin release but avoid their central adverse effects holds promise for anti-diabetic.

### Development of novel *α*_2A_AR antagonists with limit access to the central nerve system

Giving the adverse side effects of yohimbine were mainly caused by its central actions, we thus aimed to minimize its entry into the brain by structure modification. The ideal candidates should: 1) have similar or better α_2A_AR antagonistic activity as yohimbine; 2) hardly cross the blood-brain barrier (BBB); 3) have appropriate drug levels after peripheral administration. To this end, we first conducted a molecular docking analysis of α_2A_AR (PDB ID: 6KUX) with yohimbine in Schrödinger (**Fig.1a**). Our results found that the binding pocket between yohimbine and α_2A_AR is composed by H-bonding interaction, salt bridge interaction, pi-pi stacking interaction, and pi-cation interaction. These observations suggested that the aromatic phenylethylamine structure in the yohimbine is crucial for its binding and antagonistic activity to the α_2A_AR. Moreover, the binding pocket between yohimbine and α_2A_AR is narrow, while the ester group of the tail space is open to solvent, which is suitable for further modification **(Extended Data Fig.1**). Specifically, the ester group was first replaced with oxadiazole by using bioisosteric strategy, then the oxadiazole group was further modified by adding different substituents on the oxadiazole ring (**Fig.1b**). Based on this strategy, we produced more than 30 different compounds with structural diversity, with 10 of them were found with relatively similar α_2A_AR antagonistic activity as yohimbine (**Fig.1c**).

**Fig. 1.**
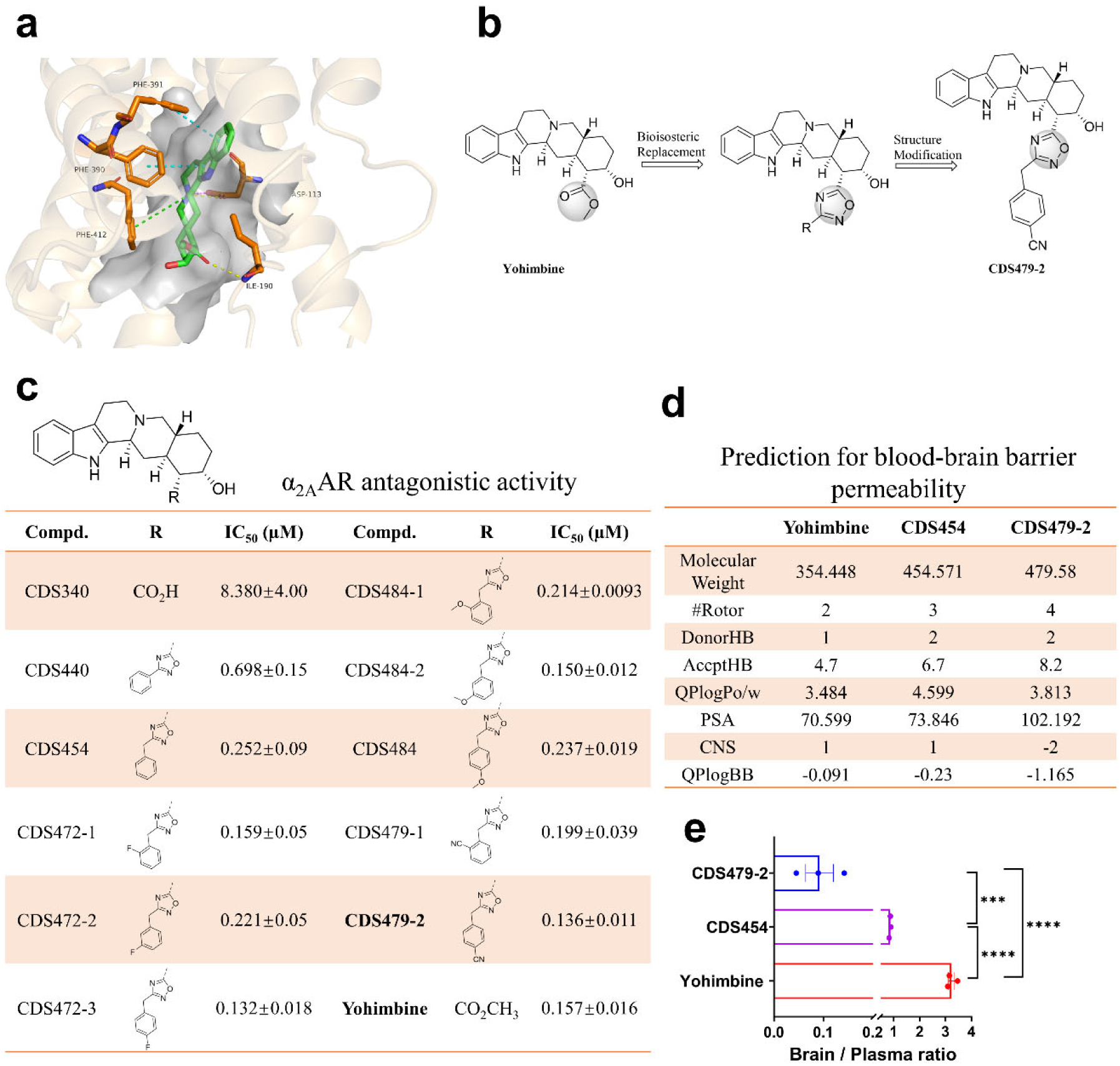
Development of novel peripherally acting adrenergic *α*_2A_AR antagonists by modification of the structure of yohimbine. **(a)** Molecular docking analysis of α_2A_AR (PDB ID: 6KUX) with Yohimbine. Yohimbine is shown in green, and adjacent amino acids are shown in orange. Yellow dashed lines indicate H-bonding interaction. Magenta dashed line indicates salt bridge interaction. Cyan dashed line indicates pi-pi stacking interaction. Green dashed line indicates pi-cation interaction. Grey section indicates ligand binding pocket. All the procedures are performed in Schrodinger, and the figure is generated by PyMOL software. **(b)** Strategy for development of novel α_2A_AR antagonists by modification of the structure of yohimbine. **(c)** *In vitro* α_2A_AR antagonistic activity test for yohimbine and the newly synthesized compounds (named after CDS). IC_50_ values were obtained from one experiment with three replicates. **(d)** Prediction of blood-brain barrier (BBB) permeability of compounds (yohimbine, CDS454, and CDS479-2) using the Qikprop of Schrödinger. The structural modification strategies to decrease the blood-brain barrier permeability include: reducing of lipophilicity (Log P), increasing of hydrogen bond donors (HBDs), molecular weight (MW), molecular polar surface area (PSA), and the number of rotatable bonds (#Rotor), etc. Lower values of CNS and QPlogBB predict lower permeability to the BBB. **(e)** The brain/plasma ratio for yohimbine (3.23±0.11), CDS454 (0.86±0.02), and CDS479-2 (0.09±0.03) in normal C57BL/6J mice at 2h after a single dose of acute administration in mice (10mg/kg, gavage, n=3 animal for each group). ***, p<0.001; ****, p<0.0001; One-way ANOVA.

The BBB permeability of drugs were largely determined by a complex of their basic structure properties, such as the number of rotatable bonds, molecular weight, the number of hydrogen bond donors, the molecular polar surface area, and the lipophilicity^17,18^. To avoid the time-consuming and expensive cost for experimental measurements of BBB permeability for the compounds, we first employed Qikprop to have a quick prediction for the new synthetic compounds. The results suggested that CDS454 and CDS479-2, which are the two that have close α_2A_AR antagonistic activity as yohimbine, have decreased permeability to the BBB compared with yohimbine, with the CDS479-2 would be the lowest one (**Fig.1d**). To verify the prediction and also to determine their drug levels, we measured their concentrations in the mice tissues at 2h after acute gavage administration(10mg/kg) (**Extended Data Table1**). The results showed that yohimbine had the highest concentrations in the plasma and all the tissues including brain, heart, liver, pancreas, and kidney. For yohimbine, the average plasma concentration was 1160ng/ml, the average brain/plasma ratio was 3.22, and the average pancreas/plasma ratio was 4.60 (**Fig.1e**). These data indicated that both the brain and pancreas are the two important action sites of yohimbine. For the 2 new compounds, CDS479-2 had the highest plasma levels and the lowest brain/plasma ratio. The average plasma concentration for CDS479-2 was 240ng/ml, and the average pancreas/plasma ratio was 0.99, and its average brain/plasma concentration ratio was 0.08, indicating that CDS479-2 indeed had the lowest permeability to the BBB (**Fig.1e**). Thus, these results suggest that the values of CNS and QPlogBB in the compounds can reliably predict their permeability to the BBB. Importantly, based on our modification strategy, we have successfully developed a novel α_2A_AR antagonist that hardly crosses the BBB.

### Cryo-EM structure of Yohimbine-bound *α*_2A_AR and CDS479-2-bound *α*_2A_AR

To validate the docking model, we elucidated the structure of the yohimbine-α_2A_AR complexes at resolutions of 3.51 Å utilizing single-particle cryo-electron microscopy (cryo-EM) (**Fig. 2a;Extended Fig.2**). High-quality electron densities of the ligands, the orthosteric binding pocket, and the transmembrane helices (TMs) were observed. Two ligands occupied the orthosteric binding pocket through a combination of broad polar, hydrophobic, and pi-pi interactions. The polar end of yohimbine formed hydrogen bonds with S215^5.42^, S219^5.46^, and Y409^6.55^, while the nitrogen atom,shared by two rings, interacted ionically with D128^3.32^. Additionally, the hydrophobic ring moieties were surrounded by hydrophobic residues, including L125^3.29^, V129^3.33^, I205^ECL2^, W402^6.48^, F405^6.51^, F406^6.52^, F427^7.39^, and F431^7.43^, among which aromatic side chains formed a pi-pi interaction network with yohimbine (**Fig. 2b-c**).

**Fig. 2.**
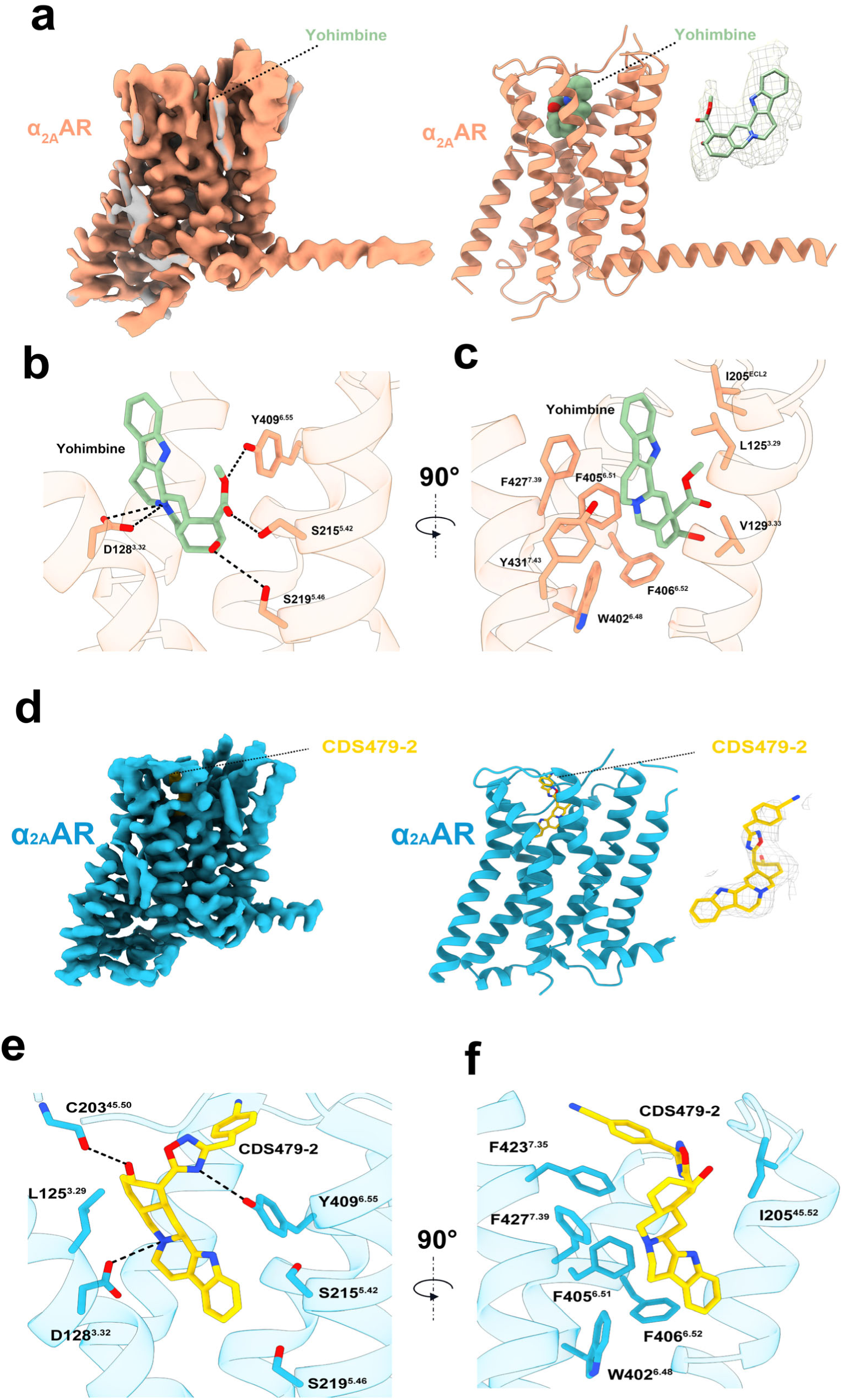
cryoEM structure of Yohimbine-bound *α*_2A_AR and CDS479-2-bound *α*_2A_AR. **a,** Electron density map (left), density of Yohimbine and 3D model (right) of Yohimbine-α_2A_AR. **b-c,** Ligand binding pocket of Yohimbine. Polar interactions are indicated by black dash lines. **d,** Electron density map (left), density of CDS479-2 and 3D model (right) of CDS479-2-α_2A_AR. **e-f,** Ligand binding pocket of CDS479-2. Polar interactions are indicated by black dash lines.

In order to compare with yohimbine and also for facilitating future structure-based optimization, we also elucidated the structure of the CDS479-2-α_2A_AR complex, which at a nominal resolution of 3.34 Å utilizing single-particle cryo-electron microscopy (cryo-EM) (**Fig. 2d; Extended Data Fig.3**). We found that the pentacyclic portion of CDS interacts with α_2A_AR mainly through van der Waals and aromatic interactions to transmembrane helices (TM) 3, 5, 6, and 7. It also forms an ionic interaction with D128^3.32^ (**Fig. 2e**), which is conserved in the other reported α_2A_AR structures^1,2^. The predicted docked pose superimposes on the cryo-EM of CDS revealed a 1.96 Å all-atom root mean square deviation (RMSD) for the antagonist (**Fig. 2f**). Consistent with the experimental result, the basic, formally cationic nitrogen of CDS is oriented towards D128^3.32^, establishing an ionic interaction (**Fig. 2f**). These results indicate the key role of D128^3.32^ in ligand binding. CDS lost the polar interactions with S215^5.42^ and S219^5.46^, compare to yohimbine-bound α_2A_AR. However, it forms a new interaction with C203^45.50^ in ECL2.

### Acute glycemic effects of yohimbine and CDS479-2 in the normal and diabetic mice

Next, we investigated the acute glycemic effects of CDS479-2 in mice, and compared with the effects of yohimbine. In the normal chow-fed C57BL/6J mice, we found that a single *i.p* injection of yohimbine or CDS479-2 dose-dependently lowered the blood glucose levels in these mice, without causing critical hypoglycemia (**Extended Data Fig.4**). The maximal glucose lowering effects of both were achieved around 30mins after acute injection and lasted for at least 2 hours (**Extended Data Fig.4**). Compared with yohimbine (1.5mg/kg), a higher dose of CDS479-2 (10mg/kg) was required to achieve similar glucose lowering effects, probably due to a relatively lower plasma and pancreas concentration of CDS479-2 after injection (**Fig.3a; Extended Data Table1**). Moreover, both CDS479-2 and yohimbine elevated blood insulin levels at 15mins after injections, but not at 30mins (**Fig.3b-c**). Interestingly, CDS479-2 treatment resulted in significantly higher insulin levels compared with yohimbine treatment at 15mins after injections (**Fig.3b**). Furthermore, pretreatment of either CDS479-2 or yohimbine significantly improved glucose tolerance **(Fig.3d-e)**. Importantly, CDS479-2 failed to lower blood glucose levels in the ADRA2A knock-out mice, indicating that α_2A_AR is necessary for its glycemic effects **(Fig.3f-g)**. Both yohimbine and CDS479-2 were found increased Ca^2+^ oscillation in the β cells of the pancreatic islets (**Extended Data Fig.5**). Given the important role of Ca^2+^ oscillation in triggering insulin vesicles release, the insulinotropic effects as observed probably mediated by direct antagonism of α_2A_AR in the pancreatic β cells. Indeed, compared to saline treatment, acute treatment of yohimbine or CDS479-2 did not significantly affect the blood glucose levels in the STZ-induced diabetic mice, in which pancreatic β cells were destructive **(Fig.3h-i)**. In the high fat diet induced obesity mice (DIO mice), again, acute injection of yohimbine or CDS479-2 significantly lowered their basal blood glucose levels (**Fig.3j)**. Pretreatment with yohimbine or CDS479-2 either by *i.p* injection or gavage administration also improved glucose tolerance in these mice (**Fig.3k-n)**.

**Fig. 3.**
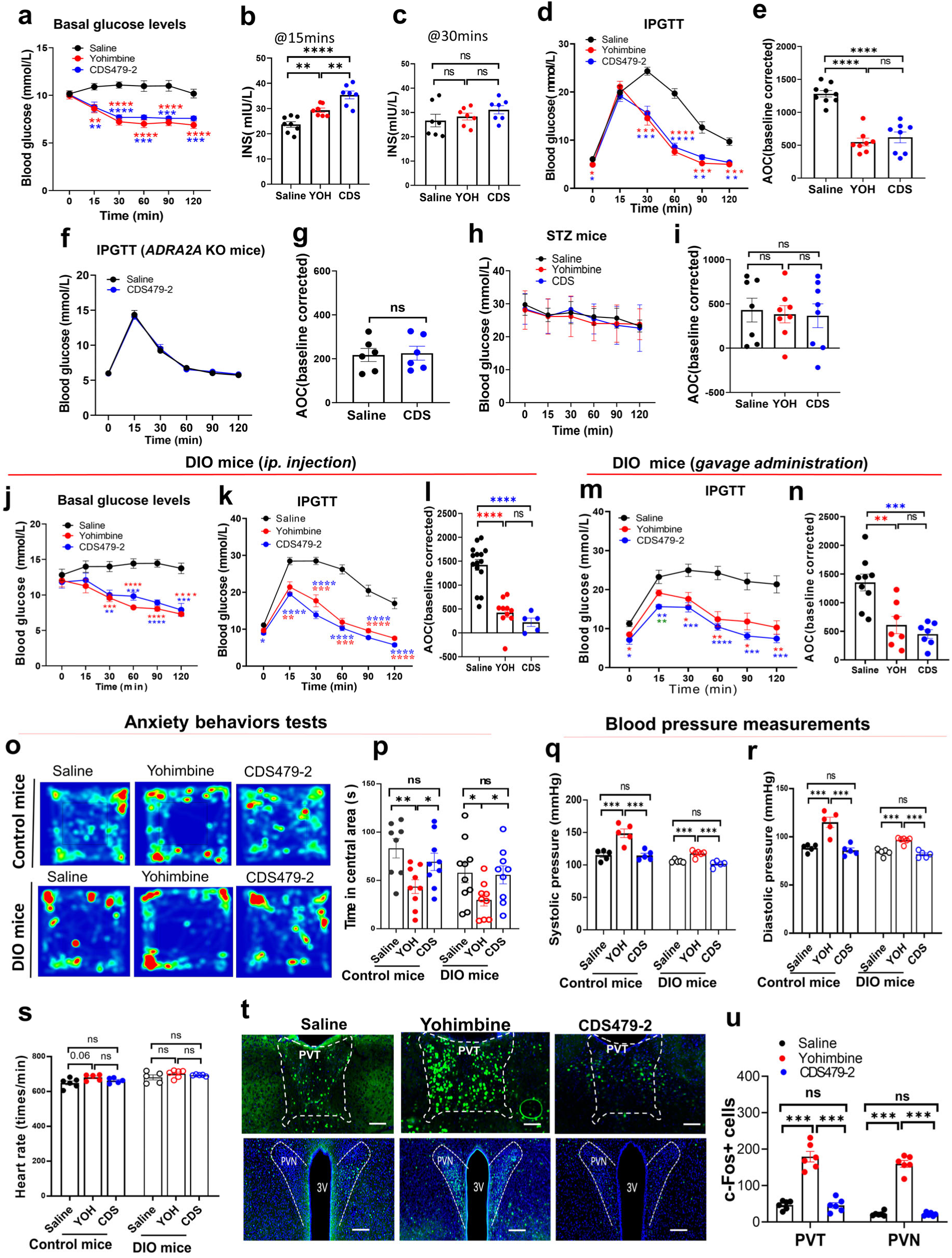
Acute administration of CDS479-2 lowers blood glucose levels, without affecting anxiety levels and blood pressures in normal and DIO C57BL/6J mice. **a-c,** Acute effect of yohimbine (1.5mg/kg, *ip*) and CDS479-2 (10mg/kg, *ip*) on basal glucose levels **(a)** and serum insulin (INS) levels **(b-c)** in normal chow-fed C57BL/6J mice. **d-g,** IPGTT (i.p glucose tolerance test) in normal C57BL/6J mice **(d)** or *ADRA2A* KO mice **(f)**. The AOC (area of curve) for GTT curve is showed in **e** and **g**, respectively. 30mins prior the test, the mice were pretreated with acute injection of saline, or yohimbine or CDS479-2. **h-i,** Acute effect of yohimbine and CDS479-2 on basal glucose levels in STZ mice (**h**).The AOC for the curve is showed in **i**. **j,** Acute effect of yohimbine and CDS479-2 on basal glucose levels of DIO C57BL/6J mice. **k-i,** IPGTT in DIO C57BL/6J mice (k). 30mins prior the test, the mice were pretreated with acute injection of saline, or yohimbine or CDS479-2. The AOC for GTT curve is showed in **l. m-n,** IPGTT in DIO C57BL/6J mice (m). 30mins prior the test, the mice were pretreated with saline, or yohimbine or CDS479-2 by gavage administration. The AOC for GTT curve is showed in **n. o-p,** Open field tests to evaluate the anxiety behaviors of normal and DIO mice after acute injection of saline, yohimbine or CDS479-2. **o,** Representative heatmaps showing the movement track of one mouse from each group. **p,** Summary data for central time. **q-s,** Blood pressures and heart rates measurements after acute injection of saline, yohimbine or CDS479-2. **q,** Systolic pressure. **r,** Diastolic pressure. **s,** heart rates. **t-u,** c-Fos expression in the PVT and PVN brain area from normal C57BL/6J mice after acute saline, yohimbine or CDS479-2 injection. **t,** representative immunofluorescence staining imaging. Green: c-Fos antibody; Blue: DAPI. scale bar: 50μm. **u,** summary data for the c-fos positive cells. *, p<0.05; **, p<0.01; ***, p<0.001,****, p<0.0001. ns: no significance. Two-way RM ANOVA with Bonferronìs post hoc test in a, d, j, k, m. One-way ANOVA with Turkey post hoc test in b, c, e, i, l, m, p, q, r, s and u. unpaired t test in g. Red asterisk: yohimbine group *vs* saline group; Blue asterisk: CDS479-2 *vs* saline group; Green asterisk: yohimbine group *vs* CDS479-2 group. Data are presented as mean ±SEM.

### Acute side effects of yohimbine and CDS479-2 in the normal and diabetic mice

To test the adverse side effects, we first performed open field test to investigate how they affected anxiety-related behaviors. We found that, after acute treatment of yohimbine (1.5mg/kg, *i.p*), both chow-fed control mice and DIO mice spent significantly less time in the central area of the open field, compared with the mice received saline or acute injection of CDS479-2 (10mg/kg, *i.p.*) (**Fig.3o-p)**. In contrast, acute treatment of CDS479-2 did not significantly affect the related behaviors in the open field test (**Fig.3o-p)**. Next, we investigated their acute effects on blood pressure by performing noninvasive tail artery blood pressure measurement. We found that a single dose of yohimbine (1.5mg/kg, *i.p*), but not CDS479-2 (10mg/kg, *i.p*), significantly increased systolic pressure (**Fig.3q**), diastolic pressure (**Fig.3r**), in both control and DIO mice. Acute treatment of yohimbine or CDS479-2 did not significantly change the heart rates in the control mice or DIO mice (**Fig.3s**). Together, these results demonstrated that acute treatment of yohimbine, at the dose that lowered blood glucose levels, also elevated anxiety levels and blood pressures in mice. These adverse effects were not observed in the CDS479-2 treatment group, probably because CDS479-2 has very low permeability to BBB (**Fig.1e**). To further confirm this idea and to identify the brain area that mediate their central actions, we performed c-Fos (a protein marker for brain activation) immunostaining on the C57BL/6J mice after acute injection of yohimbine or CDS479-2. We found that yohimbine dramatically increased c-Fos expression in the paraventricular nucleus of hypothalamus (PVN) and the paraventricular nucleus of thalamus (PVT), which brain regions that critical for mood and autonomic regulation (**Fig.3t-u**). In contrast, acute treatment of CDS479-2 did not significantly affect c-Fos expression in these brain areas.

### Chronic glycemic effects of CDS479-2 and yohimbine in the DIO mice

The pronounced glucose-lowering efficacy of a single injection of CDS479-2, without causing adverse side effects such as anxiety and elevating blood pressure, prompted us to investigate its effects of serial injections. 40-week-old DIO mice, which had been fed with 60% high-fat diet for at least 30 weeks, were then injected with CDS479-2 (10mg/kg, *i.p.*) daily for 14 days, while the aged matched DIO and chow-fed control mice were daily injected with saline for comparation (**Fig.4a**). We found that DIO mice received CDS479-2 treatment exhibited sustained glucose lowering with no significant changes in body weight, liver weight, muscle weight, fat tissues weight, food intake or anxiety levels (**Fig.4b; Extended Data Fig.6**). 3-days of daily CDS479-2 injection was sufficient to attain normoglycaemia in the DIO mice, and the effect remained stable in the subsequent injection days (**Fig.4c-d**). Furthermore, CDS479-2-treated DIO mice showed a marked improvement of glucose tolerance as revealed by the GTT performed at 48 hours after the last injection (**Fig.4e**). In the insulin sensitivity test (ITT) performed at 96 hours after the last injection, the fasting glucose levels in the CDS479-2-treated DIO mice were found still lower than the saline-treatment DIO mice, and remained lower throughout the test **(Fig.4f).** However, the insulin sensitivity was not significantly improved by the 2-weeks treatment of CDS479-2 in the DIO mice, as demonstrated in the analysis of AOC of the ITT curves **(Fig.4f inset)**. Importantly, 6 days after the last treatment, the basal glucose levels of the CDS479-2-treated DIO mice were still comparable to the control mice, and significantly lower than that of the saline-treated DIO mice (**Fig.4g)**. Consistently, the CDS479-2-treated DIO mice had significant higher plasma insulin levels than that of the saline-treated DIO mice (**Fig.4h**). The remarkable antidiabetic effects as observed above prompted us to investigate whether chronic yohimbine would have similar effects. With the same experimental procedures, we found that the DIO mice that received 2 weeks of daily yohimbine treatments (1.5mg/kg*, i.p*) did not show significant changes in the random-blood glucose levels during the treatments, compared with the DIO mice received chronic saline treatments **(Extended Data Fig.7a-d)**. Moreover, their glucose tolerance, glucose sensitivity, and the basal glucose levels after the chronic treatments, were not significantly different (**Extended Data Fig.7e-g**).

**Fig. 4.**
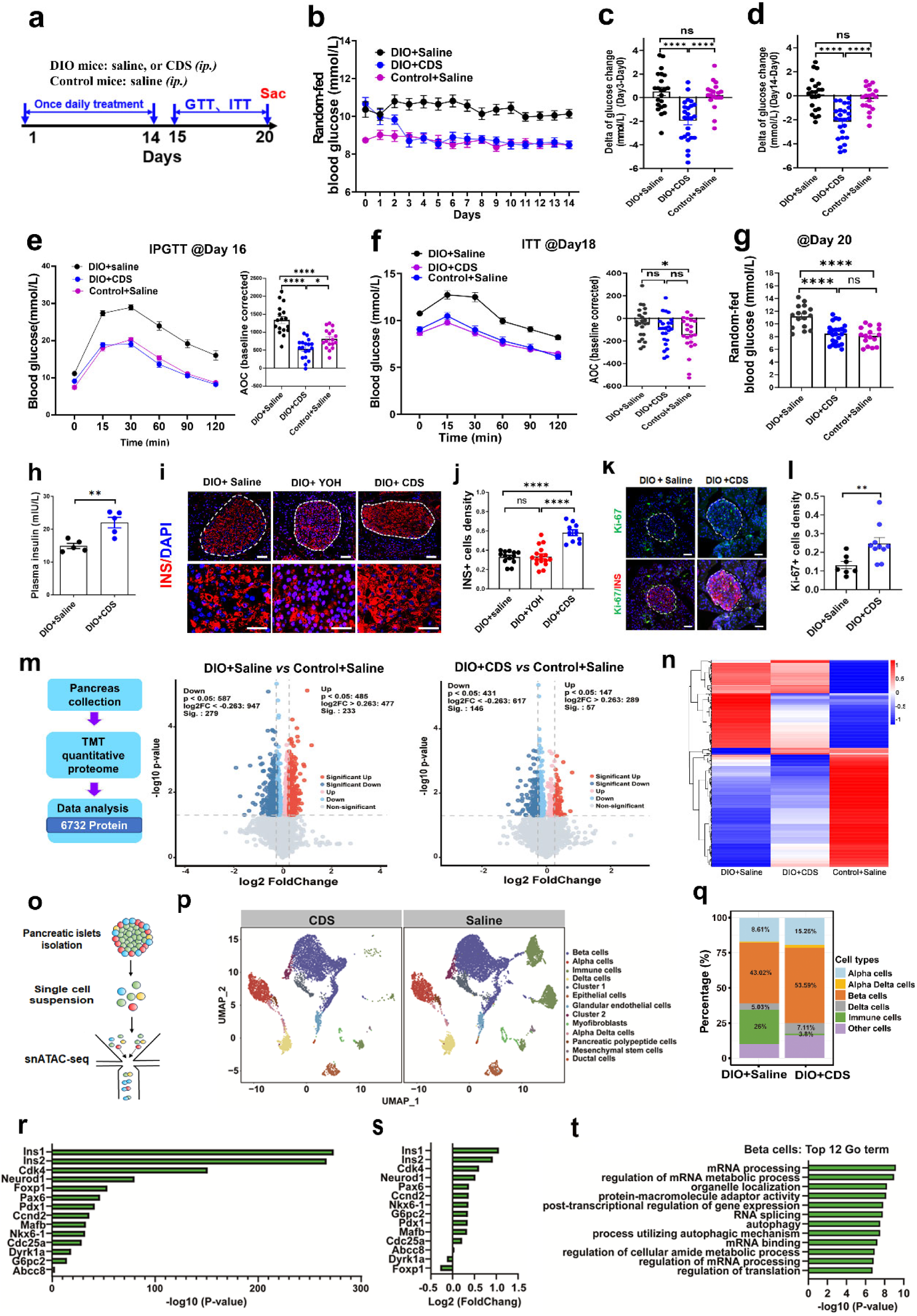
Two weeks administration of CDS479-2 achieves sustained glucose lowering effects and increases pancreatic beta cells intensity in the DIO mice. **a,** Experimental time line. The DIO mice were daily i.p. injected with saline, or 10mg/kg CDS479-2 for 14 days. Control mice were injected with saline. **b**, Daily random-fed blood glucose levels before each daily injection. **c-d**, change of random-fed blood glucose from baseline (Day 0) after 3 days of treatment (**c**), and after 14 days of treatment (**d**). **e,** IPGTT at Day 16. The inset shows the AOC of the GTT curve. **f,** ITT at Day 18. The inset shows the AOC of the ITT curve. **g,** Random-fed blood glucose levels of mice at Day 20. **h,** Plasma insulin levels of mice at Day 20. **i-j,** Insulin antibody immunostaining in the pancreatic tissues from the mice sacrificed at Day 20. **i,** A single islet from each group is delineated (up), and the magnification views of the staining are shown in the bottom. Blue, immunostaining for DAPI. Scale bar:100μm. **j,** Summary data showing the insulin positive cells density in the islets. **k-l,** Insulin and ki-67 antibodies co-immunostaining in the pancreatic islet (**k**). **l,** Summary data showing the ki-67 positive cells density in the islets. **m-n,** Proteomics analysis with the pancreatic tissues from the chronic treated mice. Volcanic maps **(m**) and heatmaps (**n**) for the differentially expressing proteins in the pancreatic tissues between groups. **o-t,** Single-cell transcriptomic analysis on the isolated pancreatic islets from the DIO mice after 2 weeks of CDS479-2 or saline treatments. **p,** UMAP analysis reveals unsupervised clustering of pancreatic islet cells in each group. **q,** The proportion of major cell types in the pancreatic islets from each group. **r-s,** Some of the statistically significant DEGs (differentially expressed genes) in the beta cells of CDS479-2 mice (**r**), and their fold changes (**s**), compared with saline-treated mice. **t,** Pathway enrichment analysis of DEGs in the beta cells of the CDS479-2 mice. Data are presented as mean values±SEM. Statistical analysis was performed with One-way ANOVA with Turkey post hoc test. *, p<0.05; **, p<0.005; ***, p<0.001. ns: no significance.

### Chronic effects of CDS479-2 on pancreatic tissues and beta cells of DIO mice at the cellular and molecular levels

Next, these chronic treated mice were sacrificed and their pancreatic tissues were harvested for biochemical, proteomic and single-cell transcriptomic analysis. First, with insulin antibodies immunostaining, we found that the density of insulin positive cells was significantly higher in the pancreatic islets of CDS479-2-treated DIO mice, compared with those of the yohimbine-treated and saline-treated DIO mice (**Fig.4i-j**). The density of insulin positive cells was not significant different between yohimbine-treated and saline-treated DIO mice(**Fig.4i-j**). Moreover, Ki-67 expression, a cell replication marker, was increased in the pancreatic islets of the CDS479-2-treated DIO mice (**Fig.4k-l**). These data may suggest that CDS479-2 treatment enhanced proliferation and regeneration of β cells in the DIO mice, which resulted in normal insulin levels and normoglycaemia in these mice. With TMT quantitative proteomic analysis of whole pancreatic tissues, we found a total of 512 differentially expressed proteins between saline-treated DIO mice and Saline-treated control mice, whereas this number narrowed down to 203 proteins in CDS479-2-treated DIO mice (**Fig. 4m)**. Thus, CDS479-2 treatment rendered the pancreas of the DIO mice more like healthy control mice in the protein expression levels **(Fig.4n)**. Q-PCR analysis with isolated pancreatic islets found that multiple genes related with proliferation and β cell were significantly upregulated in the CDS479-2-treated DIO mice, including *Pdx1, Mafa, Mafab, Neurod1,* and *Foxa2* (**Extended Data Fig.8**). To identify the molecular changes in specific cell types, we then performed scRNA-seq on the isolated pancreatic islets **(Fig.4o)**. We sequenced the transcriptomes of 19,721 pancreatic islet cells (Saline-treated DIO mice, 12,453 cells; CDS479-2-treated mice, 7,268 cells; pooled from 3-4 mice in each group). Unsupervised graph-based clustering and marker gene expression identified 12 cell types in these islets, with majority of them were endocrine cells as expected **(Fig.4p; Extended data Fig.9)**. Interestingly, a large portion of immune cells was found in the islets of the saline-treated DIO mice, which were identified by the expression of the known marker genes including *CD74, Ptrcp, Arhgap15* and *Ifi27l2a* **(Fig.4q; Extended data Fig.9)**. This observation indicated a severe inflammatory environment presented within the islets in these DIO mice, which had been under high-fat diet for at least 30weeks. Remarkably, the portion of immune cells (6%) was much less in the islets of the CDS479-2-treated mice. Importantly, the percentage of beta cells in the total islet cells was 25% higher in the CDS479-2-treated mice compared with that in the saline-treated DIO mice (53.6% vs 43.0%) **(Fig.4q)**. Moreover, the percentage of alpha cells was also higher in the CDS479-2-treated mice (8.8% vs 15.2%) **(Fig.4q)**. Further differentially expressed genes (DEGs) analysis on the beta cell population, revealed that a number of key genes/ transcription factors that related to the identity, proliferation and survival of beta cells were upregulated in the CDS479-2-treated mice, including *Ins1, Ins2, Pdx1, Pax6, Ccnd2, Mafab, Nkx6-1, G6pc2, cdc25a* etc **(Fig.4r-s)**. Pathway enrichment analysis of DEGs indicated that the top 12 pathways were mostly related to mRNA processing and their regulations **(Fig.4t)**.

## Discussion

Studies from clinical and preclinical aspects consistently indicated that α_2A_AR on pancreatic beta cells contributed to the development and pathophysiology of type 2 diabetes^1,2,7,19,20^. Pharmacologically blockage of α_2A_AR with its antagonists were found elevated plasma insulin levels and shown promising therapeutic effects for the treatment of type 2 diabetes, especially for those patients that with congenitally higher level of α_2A_AR activation of beta cells^7,21,22^. However, the clinical use of these general α_2A_AR antagonists (*ie.*yohimbine) were hampered by the adverse side effects mediated by their blockage of the α_2A_AR expressed in the central nervous system (CNS)^7^.

In this study, based on the molecular docking analysis, with bioisosteric replacement strategy on the structure of yohimbine, we have successfully developed a a novel peripherally acting α_2A_AR antagonist (CDS479-2) that has potential diabetic remission effects. Compared to yohimbine, CDS479-2 has similar α_2A_AR antagonist activity and glucose-lowering effects, but with very limited access to the CNS, and thus avoiding the unwanted side effects mediated by the antagonism of central α_2A_AR.

Strikingly, DIO mice received 2 weeks of daily CDS479-2 treatments exhibited sustained normoglycaemia and increased of beta cells density, but not the mice received yohimbine treatments. Consistent with these observations, single cell transcriptomes analysis revealed an increase portion of beta cells in the pancreatic islets from those CDS479-2-treated mice, accompanied with upregulations of a number of genes/transcription factors that critical for beta cell maturation and proliferation. Taken together, our study has uncovered that peripherally blockage of α_2A_AR promotes beta cells regeneration and restores normalization of glucose metabolism in type 2 diabetes. To the best of our knowledge, our study reveals for the first time that α_2A_AR antagonist in stimulating beta cell regeneration and sustained normoglycaemia in the diabetic animal models. Our findings also suggested that the regenerative property of beta cells in the DIO mice was constantly inhibited by the endogenous activation of α_2A_AR in these cells, probably via the tonic activity of the sympathetic innervation^23,24^. Indeed, studies with isolated pancreatic islets of rodents found that α_2A_AR agonist or norepinephrine impaired beta cell replication, which effects were blocked by α_2A_AR antagonist^25,26^. Consistently, *ADRA2A* knock-out mice exhibited increase of neonatal beta cell replication^26^. As one of the most expressed GαI-coupling GPCR in the beta cells, the activity of α_2A_AR in regulation of beta cells replication was probably through controlling the intracellular levels of cAMP, which signaling activation was found enhanced the expressions of several cell cycle-promoting proteins that essential for beta cell expansion^25,27–30^. However, these mechanic insights were largely restricted to in vitro preparations such as isolated pancreatic islets and beta cells. Moreover, given a dramatic reduction of immune cells in the pancreatic islets were revealed in the chronic CDS479-2 treated mice, whether these cells also played a role in the regeneration of beta cells are still not clear. Furthermore, the chronic effects were much less potent or absent in the yohimbine treated mice, these observations suggested that antagonism of central α_2A_AR probably inhibited beta cells regeneration via undefined mechanisms, which opposed to the effects mediated by the α_2A_AR of beta cells. Thus, future studies are required to provide more clear mechanic insights underlying the chronic effects of CDS479-2 and yohimbine in the diabetic animal models.

Our study demonstrated that the values in the Qikprop analysis for the compound structure reliably predicted the permeability to BBB. Increase of molecular weight leads to a reduction in cell membranes penetration, and decrease of lipid solubility results in poor interactions with the phospholipid bilayer, and thus limit their brain penetrations. Moreover, compounds that are highly lipid-soluble tend to have a low hydrogen bonding capacity, with each pair of hydrogen bonds reduces the permeability of the membrane by a factor of 10. Increase of PSA values was also found associated with lower BBB permeability, studies have found that compounds with PSA >90 indicated were less likely to cross the BBB. In addition, Cbrain/Cplasma and log (Cbrain/Cblood) (abbreviated as log BBB), are widely used as indicators to predict the brain distribution of the molecules, in which a Cbrain/Cplasma value of less than 0.1 is usually indicating limited brain access.

Our study demonstrated that the values from the Qikprop analysis for the compounds structure could be useful for quick prediction of their permeability to BBB. Molecules that can passively diffuse into the brain usually have small molecular weight and strong lipid solubility^31^. Compared with yohimbine, CDS479-2 has an increased molecular weight and better hydrogen bonding ability. Indeed, it was found that each pair of hydrogen bonds in the structure could indicate a reduction of the membrane permeability by a factor of 10^32^. Moreover, the Qikprop analysis found that the PSA of CDS479-2 was greater than 90 Å^2^, which is consistent with previous findings that compounds with PSA>90 Å^2^ are less likely to cross the BBB^31^. In addition, a Cbrain/Cblood value less than 0.1 usually indicates limited brain access. Again, the Qikprop analysis found that the log BBB (log (Cbrain/Cblood)) of CDS479-2 was less than −1, another evidence suggested poor BBB permeability^17^.

Overall, our study may provide a new direction for type 2 diabetes treatment. Importantly, the strategy we employed in this study to separate the beneficial and adverse side effects of yohimbine, will inspire the optimization for other drugs acting on both peripheral and central targets, such as neurotransmitters /modulators /cytokines receptors and ion channels.

## Acknowledgments

This work was supported by grants from the STI2030-Major Projects (2022ZD0207100 to ZC), the National Natural Science Foundation of China (NSFC) (31900728 to X.A.L, 32000710 to ZC, U20A2016 to ZC), the National Key Research and Development Program of China (2020YFA0908200, 2021YFA1101304 to H.L.), the Guangdong Basic and Applied Basic Research Foundation (2019A1515110190 to ZC, 2023A1515011743 to X.A.L, 2023A1515010483 to L.C.), Shenzhen Key Basic

Research Project (JCYJ20200109115641762 to ZC), the Young Scientists Program of Guangzhou Laboratory (QNPG23-02 to L.C.), Shenzhen governmental grant (ZDSYS20190902093601675 to ZC), Shenzhen Science and Technology Program (JCYJ20210324134004013 to JL).

## Authors contribution

The original idea to modify the structure of yohimbine to avoid its central effects was from Z.C. The detail research was designed and supervised by X.L, Y.D, J.N, Z.C. The animal studies, including blood glucose measurements, immunostainings, behaviors, proteomics analysis, single-cell RNA-seq analysis, and Q-PCR were done by the team of X.L and Z.C, in which F.H, Y.X, X.J, R.W, Z.X, J.R, C.S, S.J, X.H, Y.Z, X.L, Z.C conducted the experiments. The structure modification of yohimbine, tissues distribution of the compounds, molecular docking analysis, BBB permeability prediction, α_2A_AR antagonist activity tests were done by the team of J.N and X.X, in which J.W, D.C, X.Xu conducted the experiments. Th Cryo-EM data were collected and analyzed by the team of Y.D, in which Q.W, S.G, F.Y conducted the experiments. In vitro studies were done by L.C, T.Z, H.R, supervised by L.C and H.L. W.L, J.D, L.W, C.T and T.X contributed new reagents/analytical tools; F.H, J.W, D.C, Q.W, S.G, F.Y, H.R, L.C and Z.C analyzed data; F.H, J.W, Y.F, S.G and Z.C wrote the paper.

## Extended information

**Extended Table 1.**
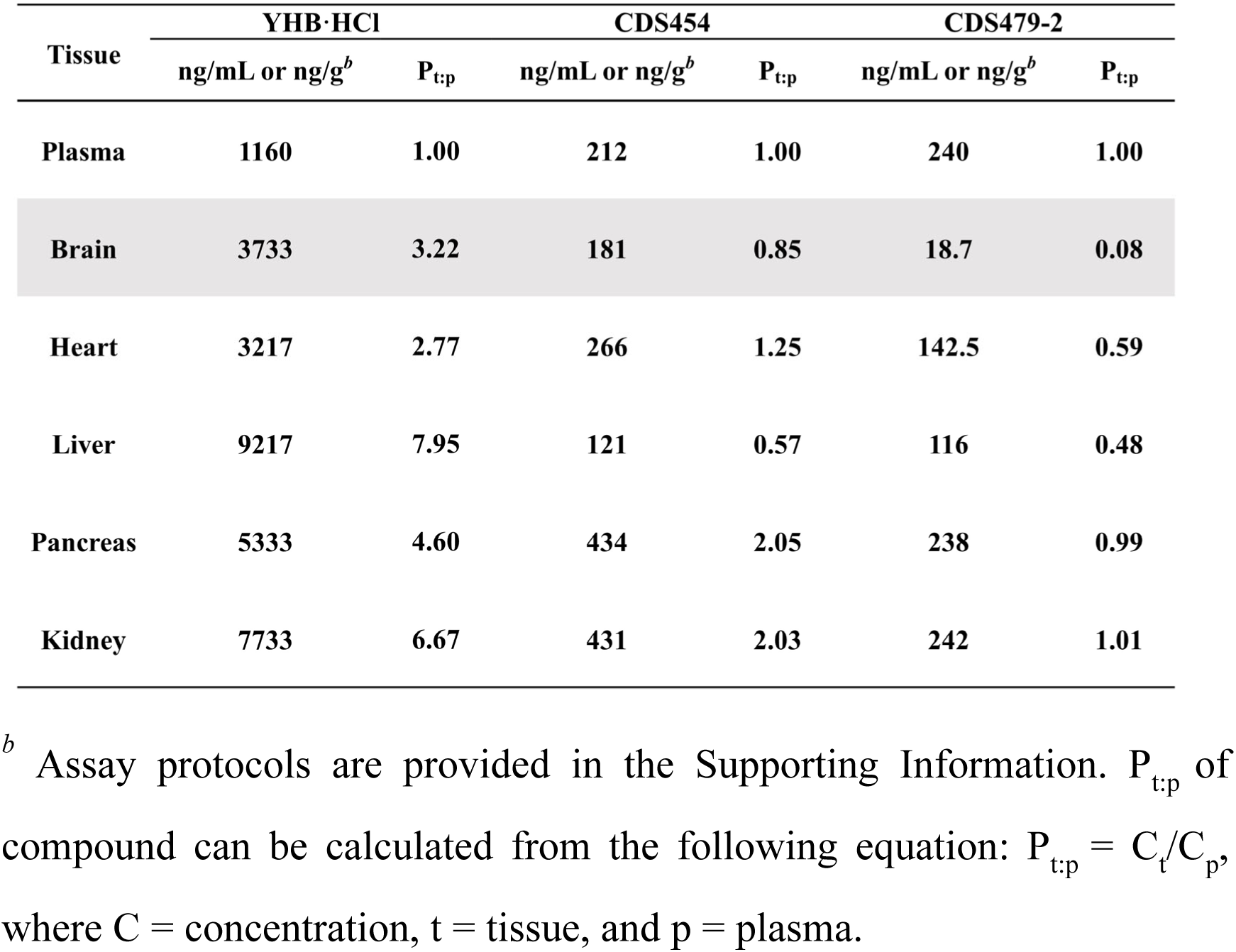
Tissue distribution of prototype in male mice 2 hours after gavage administration of YHB and yohimbine derivatives (10 mg/kg).

| Tissue | YHB·HCl |  | CDS454 |  | CDS479-2 |  |
| --- | --- | --- | --- | --- | --- | --- |
|  | ng/mL or ng/g<sup>b</sup> | P<sub>t:p</sub> | ng/mL or ng/g<sup>b</sup> | P<sub>t:p</sub> | ng/mL or ng/g<sup>b</sup> | P<sub>t:p</sub> |
| Plasma | 1160 | 1.00 | 212 | 1.00 | 240 | 1.00 |
| Brain | 3733 | 3.22 | 181 | 0.85 | 18.7 | 0.08 |
| Heart | 3217 | 2.77 | 266 | 1.25 | 142.5 | 0.59 |
| Liver | 9217 | 7.95 | 121 | 0.57 | 116 | 0.48 |
| Pancreas | 5333 | 4.60 | 434 | 2.05 | 238 | 0.99 |
| Kidney | 7733 | 6.67 | 431 | 2.03 | 242 | 1.01 |

<sup>b</sup> Assay protocols are provided in the Supporting Information. P<sub>t:p</sub> of compound can be calculated from the following equation: $P_{t:p} = C_t/C_p$ , where C = concentration, t = tissue, and p = plasma.

**Extended Data Fig.1.**
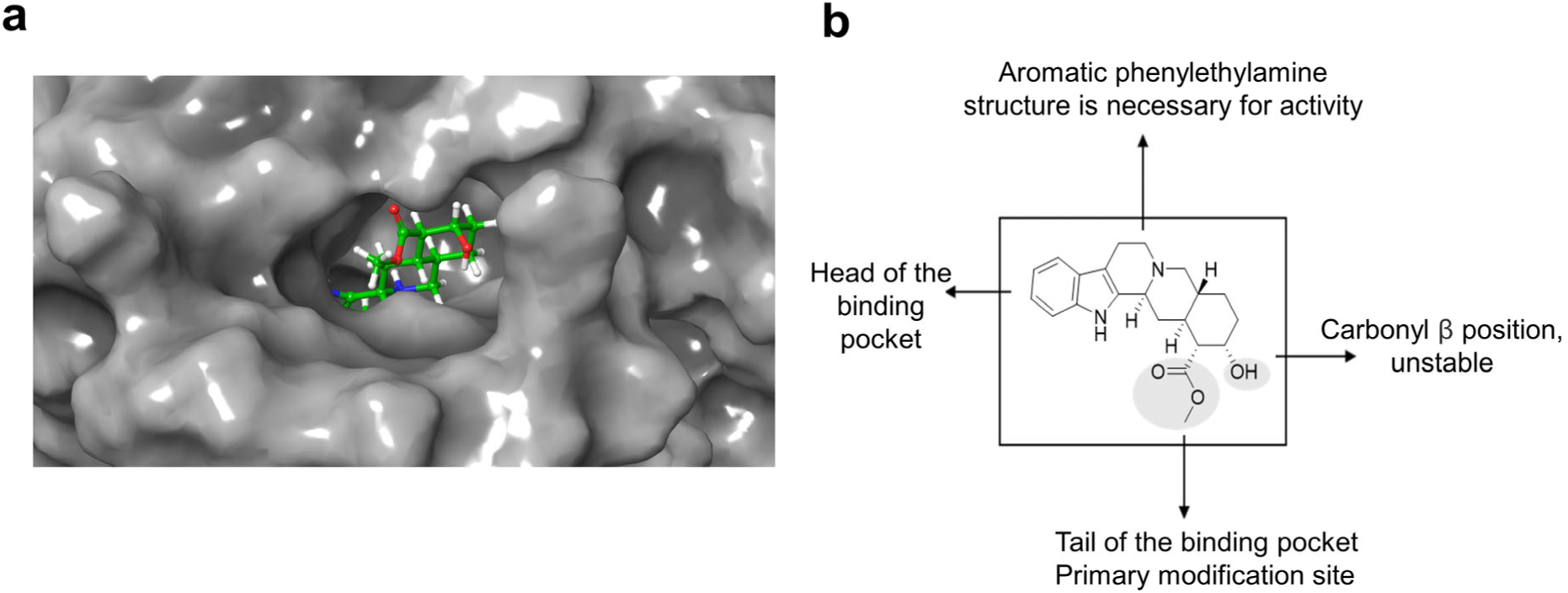
Molecular docking analysis of *α*_2A_AR (PDB ID: 6KUX) with yohimbine. **a,** The position of yohimbine in the α_2A_AR binding pocket. Yohimbine is shown in green and α_2A_AR is shown in grey. b, Analysis of the modification sites of yohimbine.

**Extended Data Fig.2.**
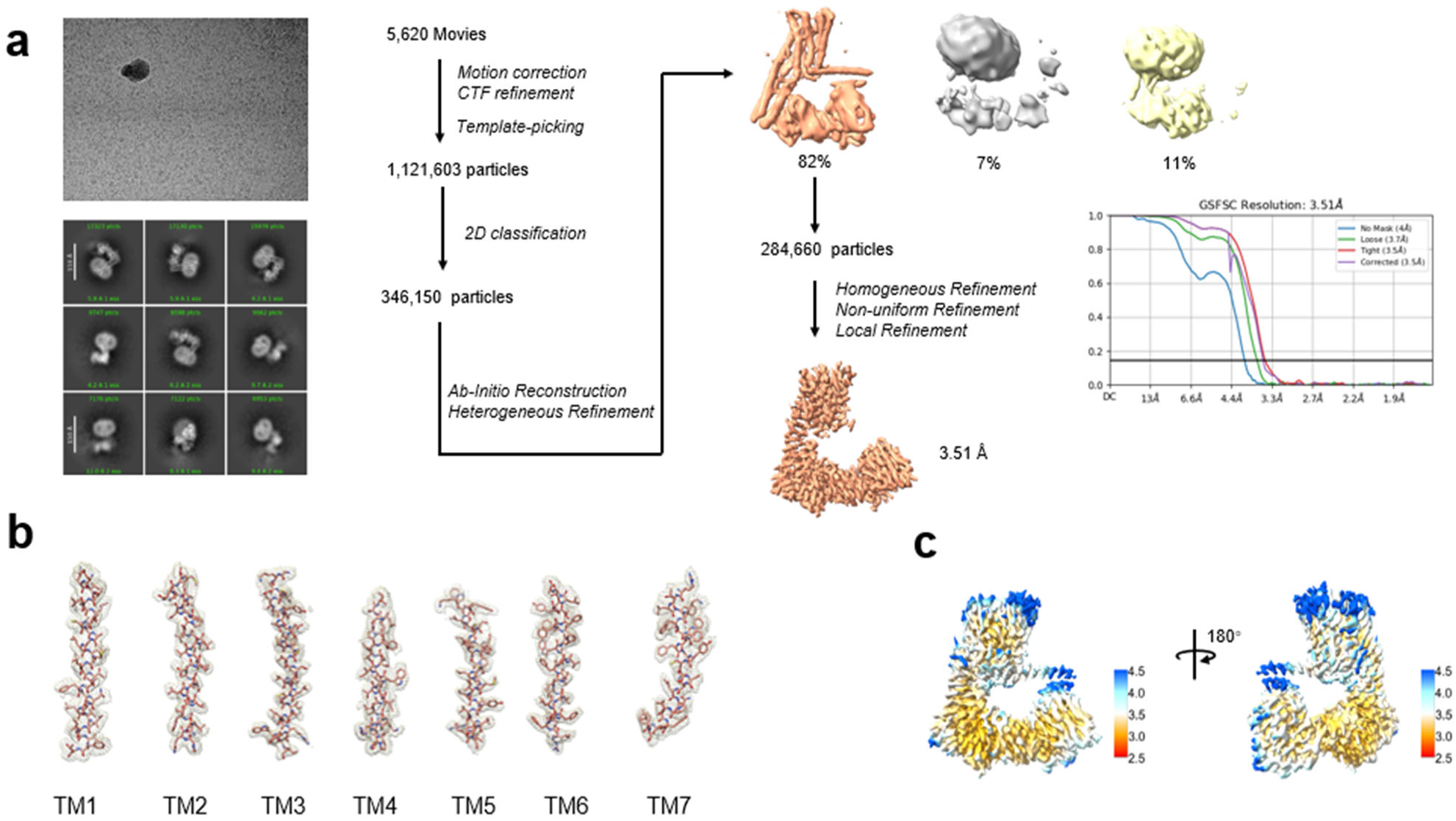
Single-particle reconstruction of YOH-bound *α*_2A_AR. **a,** Cryo-EM micrograph, 2D class averages, flow chart and FSC of cryo-EM analysis for YOH-α2AAR; **b,** Side-chain density of TM helix of α2AAR; **c,** Local resolution display of the reconstructed 3.51 Å map. A local resolution of 3.51 Å map was projected within cryoSPARC. The resolution range is depicted from 2.5 Å-4.5 Å.

**Extended Data Fig.3.**
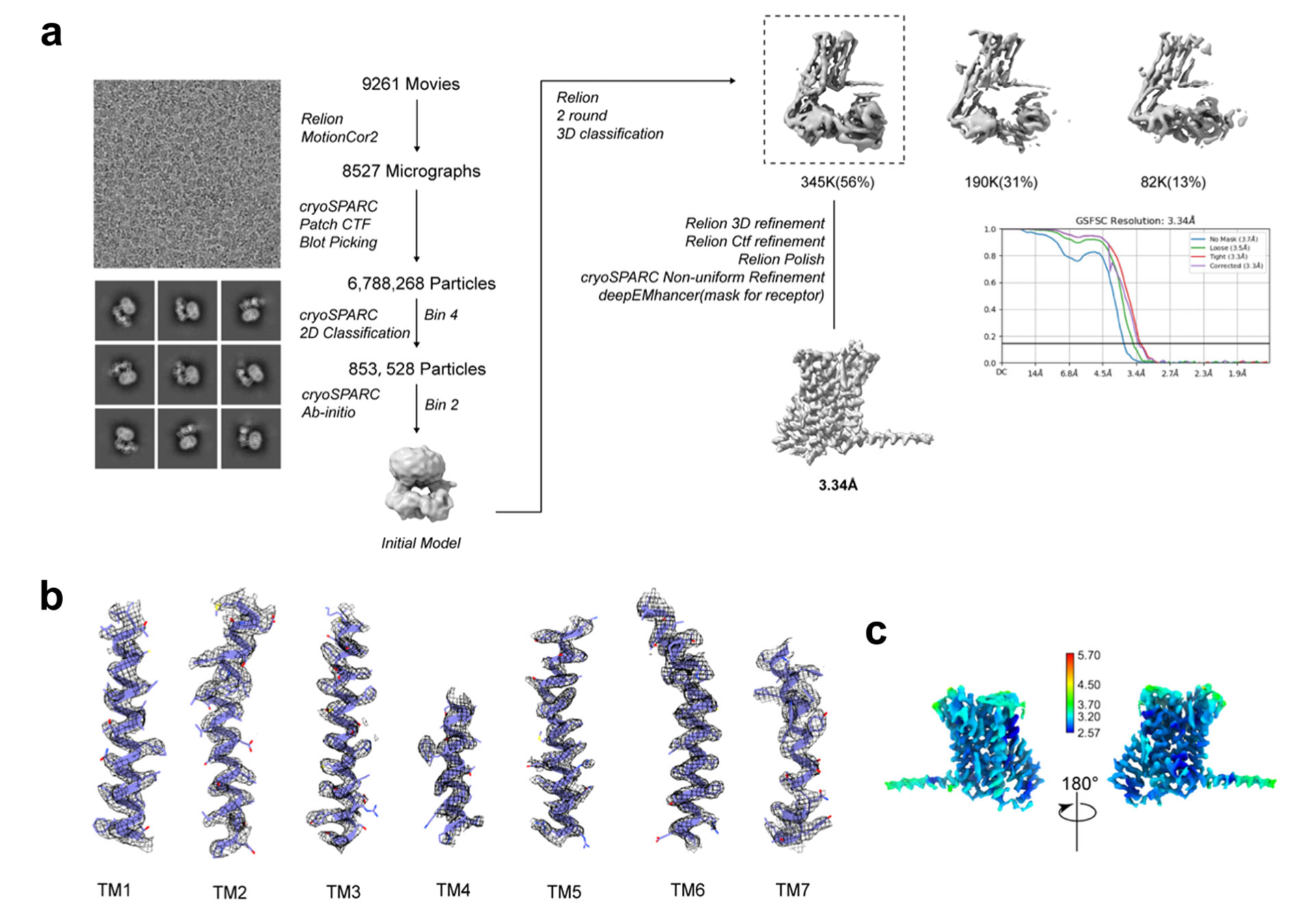
Single-particle reconstruction of CDS479-2-bound *α*_2A_AR. **a,** Cryo-EM micrograph, 2D class averages, flow chart and FSC of cryo-EM analysis for CDS-α_2A_AR; **b,** Side-chain density of TM helix of α_2A_AR; **c,** Local resolution display of the reconstructed 3.34 Å map. A local resolution of 3.34 Å map was projected within cryoSPARC. The resolution range is depicted from 2.5 Å-5.7 Å. Volume was contoured at a threshold level of 0.546.

**Extended Data Fig.4.**
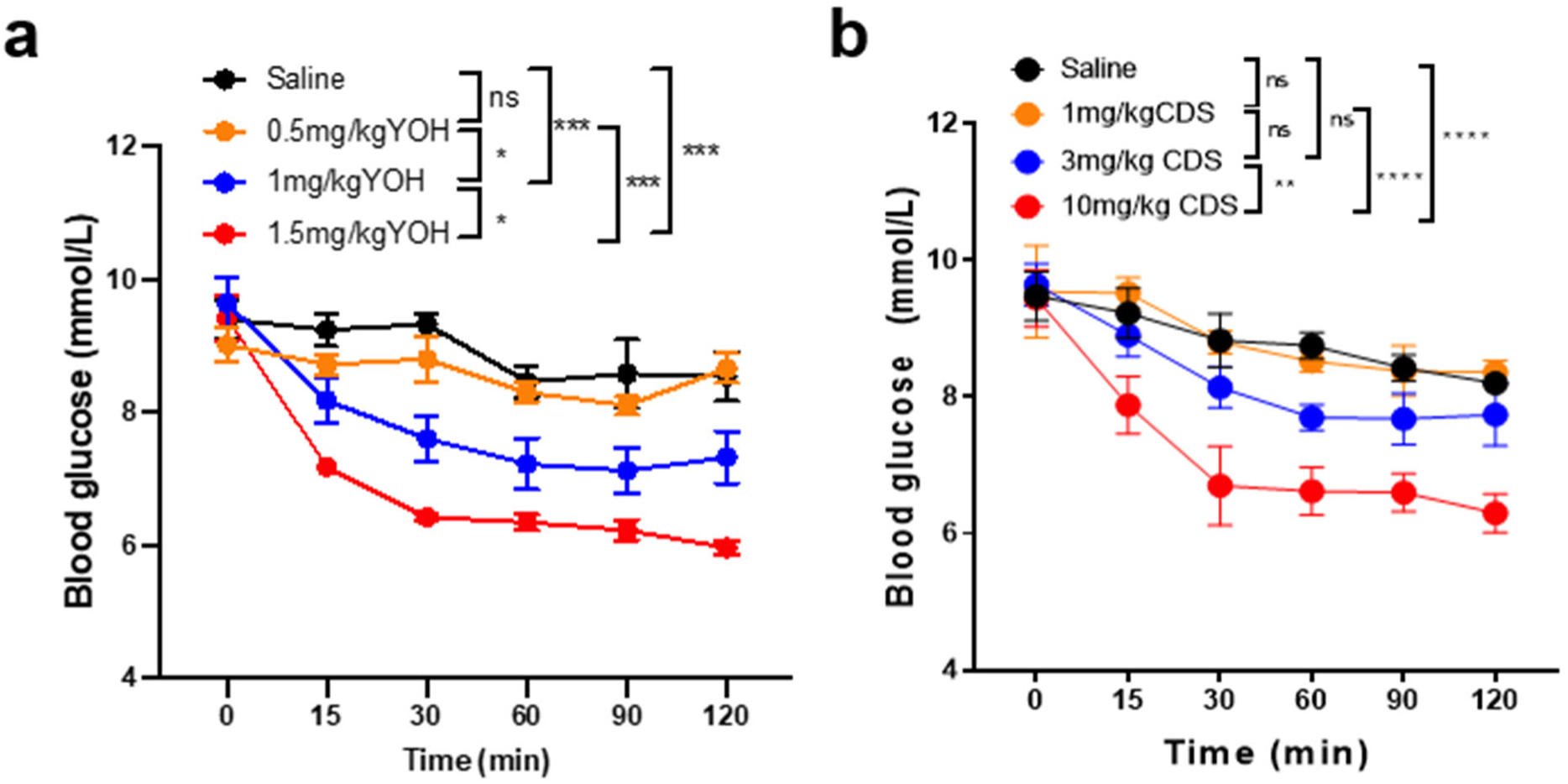
Dose-depend effects of yohimbine and CDS479-2 on basal blood glucose of normal fed C57BL/6J mice. **a,** Acute effects of different doses of yohimbine (YOH) on basal blood glucose of normal fed C57BL6J mice. **b,** Acute effects of different doses of CDS479-2(CDS) on basal blood glucose of normal fed C57BL6J mice. The drugs were injected (*i.p*) at time 0. Blood glucose levels of the mice were measured at 15, 30, 60, 90, 120 after injections. n=5 animals for each group. ns: no significance, * p<0.05, ** p<0.01, *** p<0.001, **** p<0.0001. Two-way RM ANOVA. Data are presented as mean ± SEM.

**Extended Data Fig.5.**
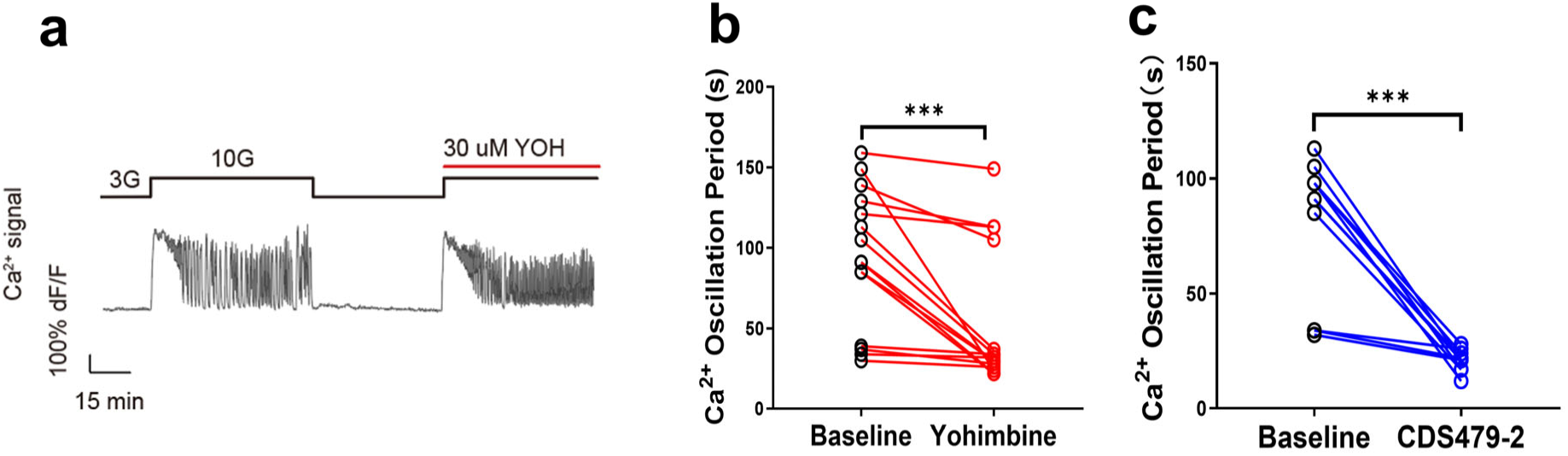
Yohimbine and CDS479-2 increased Ca^2+^ oscillation in the *β* cells of the pancreatic islets. **a,** Recordings of whole islet Ca^2+^ signal with 3G, 10G, 30 uM YOH. **b,** Oscillation period before treatment (10G), after YOH (n=15 islets from 3 *Ins1-Cre^+/-^; GCaMP6f^f/+^* mice). **c,** Oscillation period before treatment (10G), after CDS (blue; n=10 islets from 3 *Ins1-Cre^+/-^; GCaMP6f^f/+^* mice). ***,p<0.001; Paired t test.

**Extended Data Fig.6.**
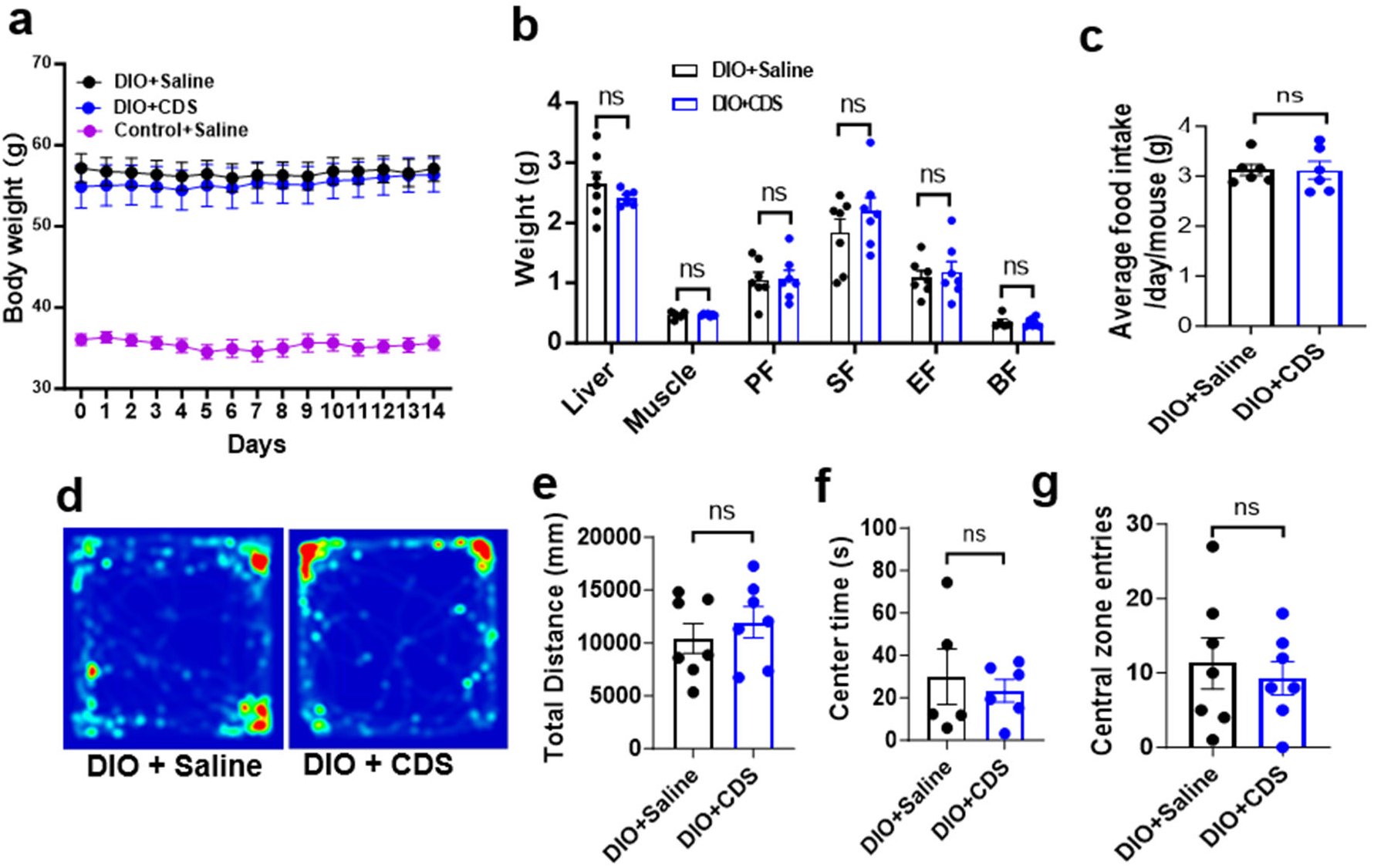
2 weeks daily treatment of yohimbine or CDS did not significantly change the body weight, food intake and anxiety levels in DIO mice. **a,** body weight. **b,** body fats. **c,** daily food intake for each mouse. **d-g,** open field test. The total distance the mice movement in the open field(e). The time spent in the central zone(f). The entries into the central zone (g). ns: no significance; unpaired t test.

**Extended Data Fig.7.**
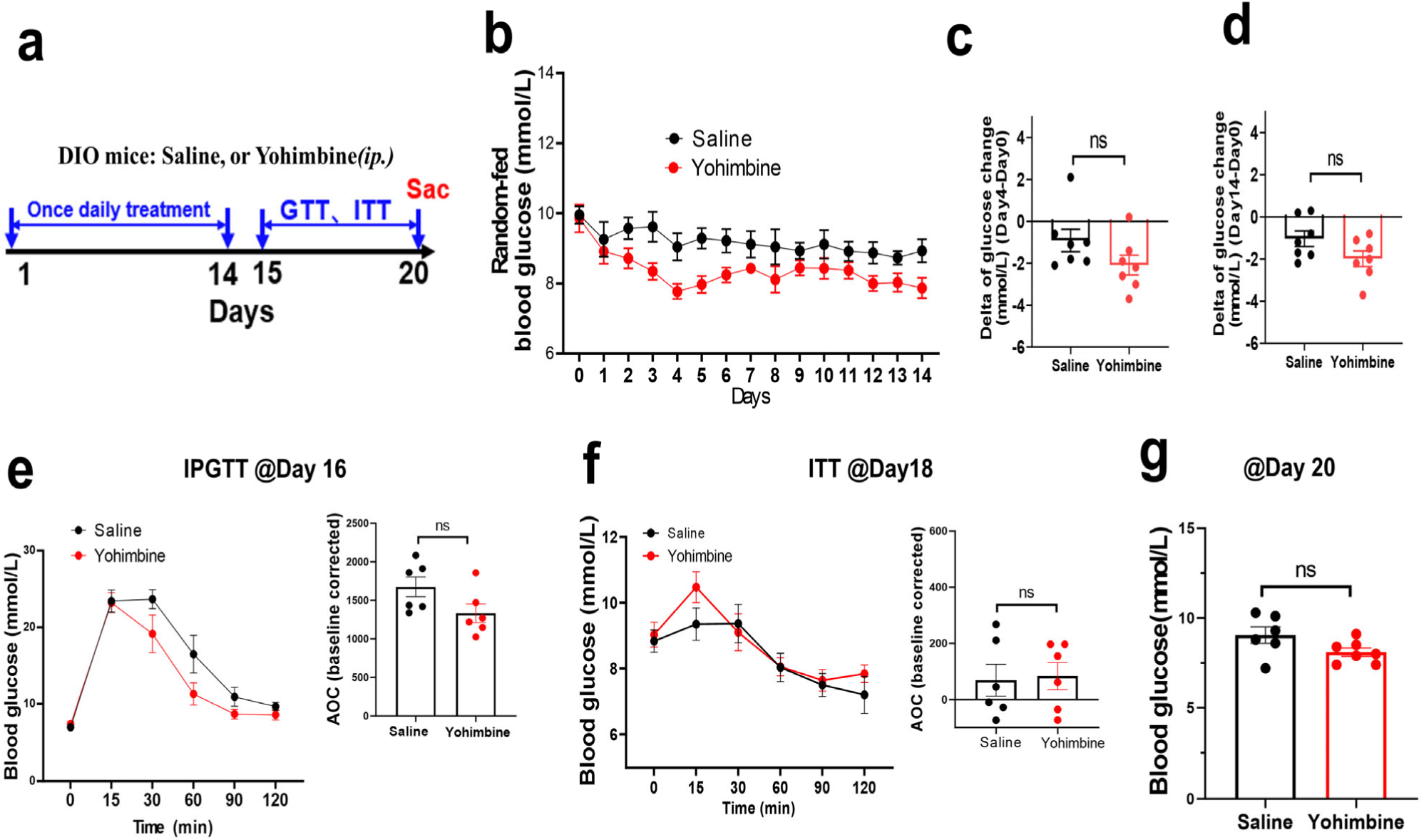
The effects of 2 weeks daily treatment of yohimbine on the blood glucose levels in the DIO mice. **a,** Experimental time line. The DIO mice were daily *i.p.* injected with saline, or 1.5mg/kg CDS479-2 for 14 days. **b**, Daily random-fed blood glucose levels of the mice during treatments. **c-d**, Change of random-fed blood glucose from baseline (Day 0) after 3 days of treatment (**c**), and after 14 days of treatment (**d**). **e,** IPGTT at Day 16. Th inset shows the AOC of the GTT curve. **d,** ITT at Day 18. The inset shows the AOC of the ITT curve. **g,** Random-fed blood glucose levels of mice at Day 20. 6 days after last treatment.

**Extended Data Fig.8.**
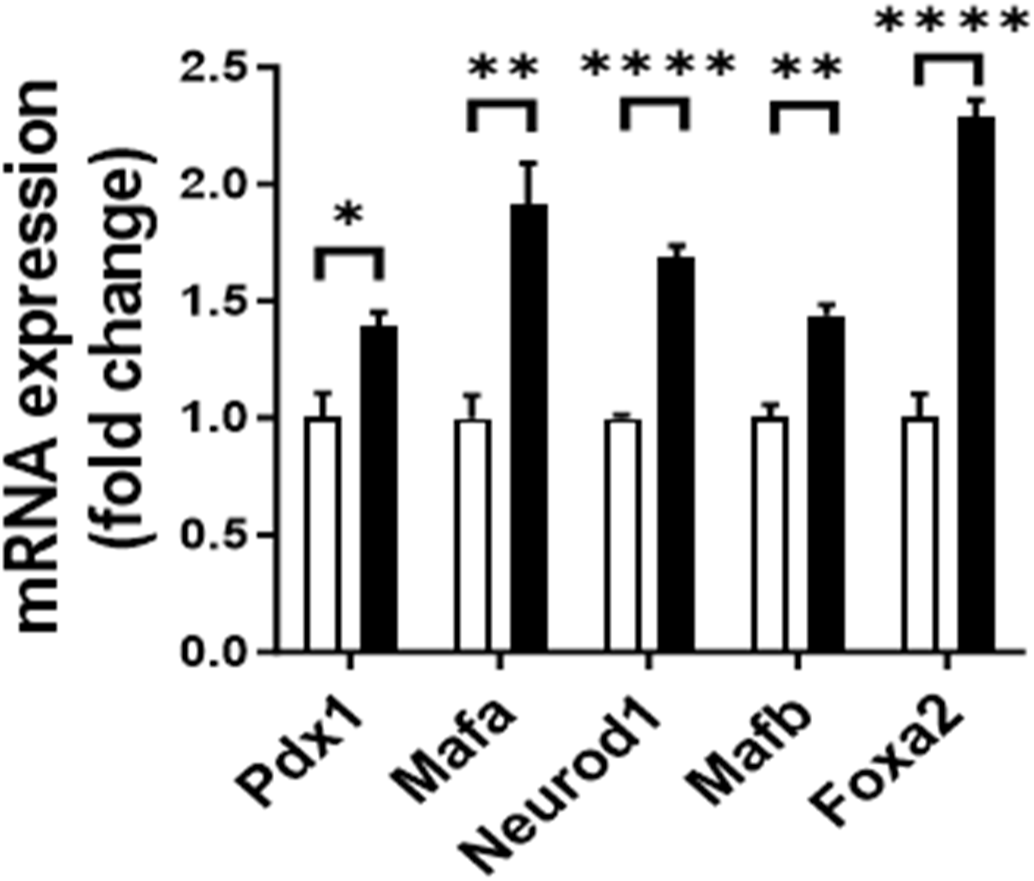
Q-PCR results found that proliferation marker genes were upregulated in the pancreatic islets of CDS479-2 treated DIO mice, compared with saline-treated DIO mice.

**Extended Data Fig.9.**
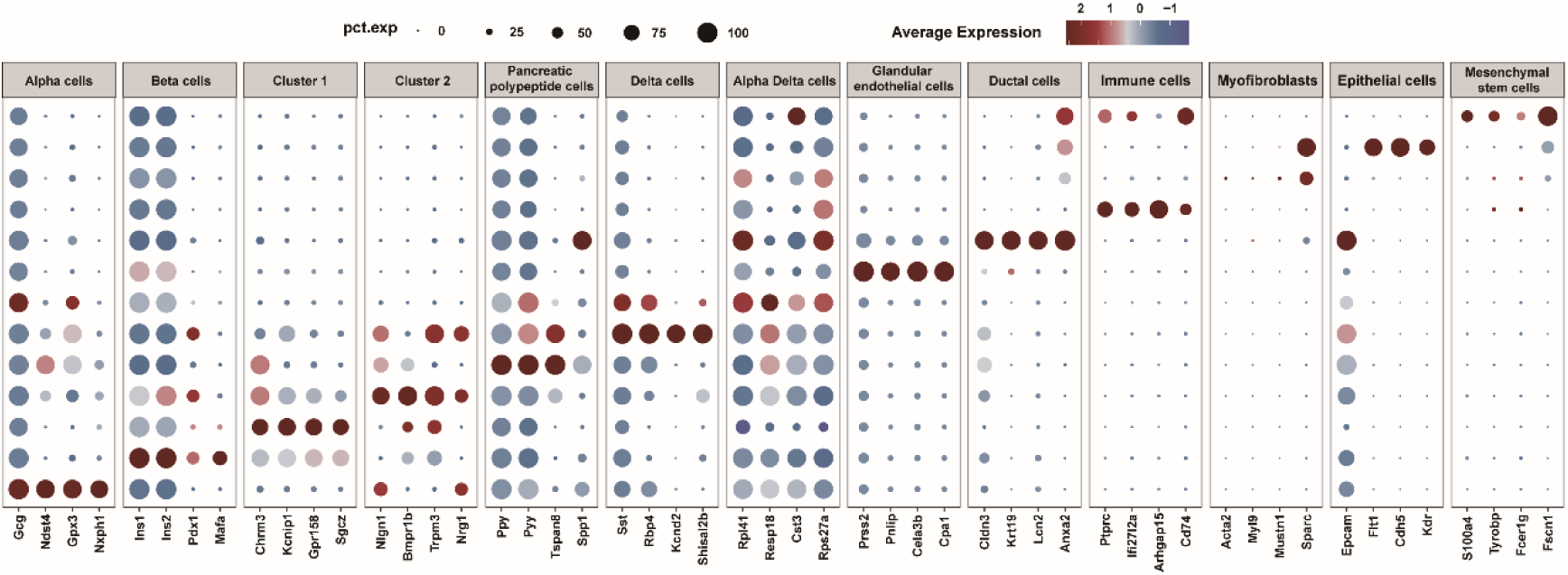
Gene markers for each cell type of the pancreatic islets revealed by unsupervised clustering with scRNA-seq analysis in the chronic CDS479-2-treated and saline-treated DIO mice.

## Materials and Methods

### Animals

All male mice were housed with 5 per cage and maintained in SPF animal facility at room temperature, 12 h light/dark cycle, and free access to food and water, unless indicated wherever possible. Mice were randomly assigned for each experiment. Behavior and glucose measurements were performed at the similar time of the day. All animal experiments were performed under the institutional guideline of IACUC of Shenzhen Institute of Advanced Technology, Chinese Academy of Sciences. Male C57bl/6J mice age between 8-16 weeks-old were used as normal healthy control mice. db/db and ob/ob mice at 8-weeks-old were purchased from Zhejiang Weitong Lihua Co. The Ins1-Cre mice and Rosa26-GCaMP6f^flox^ mice were purchased from Jackson Laboratory. For diet induced obesity (DIO) type 2 diabetic mice model, male C57bl/6J mice at 8 weeks age were fed with a 60% high-fat diet (HFD, D12492, Soest, Germany) for at least 30 weeks, while the control mice were fed with chow diet. The bodyweight, food intake and random blood glucose were monitored every week.

### Acute injection and pharmacokinetic Study

10-week-old male C57bl/6J mice were intraperitoneal or gavage administration with yohimbine, CDS479-2 (dissolved in deionized water) or saline at 1.5mg/kg and anesthetized 30 min later. The blood was collected and clotted at 4°C for 1 h. After a 12,000-rpm centrifugation at 4°C for 10 min, the blood serum was collected and stored for insulin measurement. The pancreas, heart, liver, kidney, and brain were taken immediately freezing with liquid nitrogen for yohimbine and CDS479-2 measurement. Parts of those fresh tissue were immersed in 4% paraformaldehyde for fixation. For c-Fos analysis, the mice were i.p. injected with yohimbine, CDS479-2 or saline and then anesthetized 2h later. After 4% paraformaldehyde perfusion, complete brain tissues were quickly taken from mice and immersed in 4% paraformaldehyde. For pharmacokinetic study, the samples were then analyzed by liquid chromatography tandem mass spectrometry (LC-MS/MS).

### Acute and chronic injection experiment

In acute experiments, the effects on blood glucose, blood pressure and behavioral were determined in DIO mice 30 min after a single intraperitoneal injection or gavage administration of yohimbine or CDS479-2. Two or more days were required between each experiment. In chronic experiments, DIO mice were administered yohimbine or CDS479-2 by intraperitoneal injection for 14 or 21 consecutive days, and random blood glucose and body weight were measured daily during administration. Chronic injected mice were subjected to glucose tolerance test and insulin tolerance test on days 1 and 3 after completion of the injections, respectively, and were sacrificed on day 5. The tissues were used for immunofluorescence staining experiments, pancreatic proteomic and pancreatic islet single-cell assay experiments.

### Intraperitoneal glucose tolerance test (IPGTT)

Mice were fasted for 16h (5 pm to next day 9 am, free access to water) before the test. Tail vein blood glucose levels were monitored with a Contour Plus Blood Glucose Meter (Ascensia Diabetes Care Holdings AG). Briefly, basal blood glucose levels of the mice were measured then followed with a single i.p. injection of glucose (1.5 g/kg of body weight). Blood glucose levels were measured at 15, 30 60, 90 and 120 min after glucose injection.

### Intraperitoneal insulin Tolerance Test (IPITT)

Mice were fasted for 4h (9 am to 1 pm, free access to water) before the test. Briefly, basal blood glucose levels of the mice were measured then followed with a single i.p injection of insulin (1.5 UI/kg of body weight). Blood glucose levels were measured at 15, 30 60, 90 and 120 min after insulin injection.

### Open field test

Mice were placed in a bright box (35 × 35 × 30 cm) where the animals can move freely. After placing the animal into center of the box, the movement of the mice was recorded by a video camera for 10 min. The video recording was analyzed by using Anymaze that with a floor divided into center (17.5 × 17.5 cm) and side, the time spent with moving around the box was evaluated.

### Noninvasive tail artery blood pressure measurement

The procedure was similar to a method previously described by Johns et al. (Johns C, 1996). Mice were conditioned to the mouse fixator restraint for at least 3 days before measurements. Open the ZS-Z non-invasive blood pressure measurement system of rats and mice, open the chamber and set the temperature to 33-34 °C. The mice were injected with the drugs and fixed in the mouse fixator with the abdomen facing down. Place the pulse compression sleeve and the high-sensitivity pulse transducer at the upper 1/3 of the tail in turn and fix the transducer with the surface aligned with the ventral side of the tail. After the mouse is quiet, open the Medlab biological signal acquisition and processing system, and pulse waves can be seen. If there is no typical pulse waveform, adjust the position of the sensor until the waveform appears. After the waveform was stable, the pressure in the pulse pressure cuff was increased with a balloon to allow the mice to adapt. After 30 min of administration to mice, data collection was started. Increase the pressure in the pressure cuff with a balloon until the pulse wave completely disappears, then increase the pressure by 30 mmHg, and then slowly deflate through the balloon valve. Carefully observe the pulse wave from being blocked to the first wave again. The pressure corresponding to this first wave is the systolic pressure, and the pressure corresponding to the pulse wave when it first recovers to the amplitude before blocking is the diastolic pressure (usually the fourth or fifth wave after the systolic pressure corresponds to the wave). Repeat the measurement for 5-6 times and take the average value of the stable curve. Select 5-6 segments from the typical pulse waves recorded before and 30 min after of administration to measure the heart rate and record the average value as the heart rate.

### Immunofluorescence staining

For immunofluorescence staining, sections were made on a Leica freezer slicer at - 20 ℃. brain tissue was cut into 20 μm while the pancreatic tissue was 8 μm. Sections of tissue were incubated with 5% horse serum for 30 min and then incubated with anti-insulin antibody, or anti c-Fos antibody overnight at 4 ℃. After washing with PBST, sections were incubated with goat anti mouse Alexa Fluor 488 secondary antibody, goat anti rabbit Alexa Fluor 594 secondary antibody (Cell Signaling Technology, USA) and DAPI (Sigma, USA) for 2 h. Sections were then observed under an immunofluorescence microscope.

### Serum insulin concentration measurement

The level of mouse insulin (INS) in the blood serum was determined by the double antibody sandwich method. The purified mouse insulin capture antibody was used to coat the microplate to make solid phase antibody. The mouse insulin was added to the coated microplate in turn, and then combined with the HRP-labeled detection antibody to form the antibody-antigen-enzyme-labelled antibody complex. After thorough washing, the substrate TMB was added for color development. TMB is converted into blue under the catalysis of HRP enzyme, and finally into yellow under the action of acid. The color depth is positively correlated with the mouse insulin in the sample. The absorbance (OD value) was measured with an enzyme marker at the wavelength of 450 nm, and the content of mouse insulin in the sample was calculated through the standard curve.

### Pancreas tissues preparation for proteomic analysis

Take out the pancreatic tissue sample from the - 80 ℃ refrigerator, grind it into powder at low temperature and quickly transfer it to the liquid nitrogen precooled centrifuge tube, add an appropriate amount of PASP protein lysate (100 mM ammonium bicarbonate, 8 M urea, pH=8), shake and mix well, and fully lyse it with ice water bath ultrasonic for 5 minutes. Centrifuge at 4 ℃ and 12000 g for 15 min, add 10 mM DTT to the supernatant and react at 56 ℃ for 1 h, then add sufficient IAM and react at room temperature and away from light for 1 h. Add 4 volumes of - 20 ℃ precooled acetone to precipitate at - 20 ℃ At least 2 h, centrifugate at 4 ℃ and 12000 g for 15 min, and collect the sediment. Then add 1mL - 20 ℃ precooled acetone for resuspension and wash the precipitate, centrifuge, and collect the precipitate, air dry, and add appropriate amount of protein solution (8 M urea 100 mM TEAB, pH=8.5) dissolved protein precipitation.

### Ca^2+^ Oscillation recording from the Pancreatic Islets

Islets of Langerhans were isolated from *Ins1-Cre+/-; GCaMP6f^f/f^*mice, which were generated by crossbreeding Ins1-Cre mice (Jackson Laboratory, Strain #:026801) with Rosa26-GCaMP6f^flox^ mice (Jackson Laboratory, Strain #:028865). After isolation, the islets were cultured overnight in RPMI-1640 medium containing 10% fetal bovine serum (10099141C, Gibco), 8 mM D-glucose, 100 unit/ml penicillin and 100 mg/ml streptomycin for overnight culture at 37 ℃ in a 5% CO_2_ humidified air atmosphere. The islet in the microfluidic chip was kept at 37 ℃ and 5% CO_2_ on the microscope stage during imaging. The reagents were automatically pumped into the microfluidic chip with a flow rate of 400 μL/h by the TS-1B syringe pump (LongerPump), which was controlled by software written in MATLAB. All fluorescence images were acquired using spinning-disc confocal microscopy (10x objective, Dragonfly) at time resolution of 4.5 s per frame.

### TMT-labeled proteome spectrometric analyses

After determination of protein concentration, add 100 μL 0.1 M TEAB buffer solution was re dissolved and 41 μL TMT labeling reagent dissolved in acetonitrile was mixed upside down at room temperature for 2h. After that, add 8% ammonia water with final concentration to terminate the reaction, take samples with equal volume after labeling, mix them, and freeze dry them after desalination. Prepare mobile phase solution A (2% acetonitrile, 98% water, ammonia water adjusted to pH=10) and solution B (98% acetonitrile, 2% water). Dissolve and mix the freeze-dried powder with solution A, and centrifuge 12000 g at room temperature for 10 min. Use L-3000 HPLC system, chromatographic column Water BEH C18 (4.6 × 250 mm, 5 μm), the column temperature is set at 45 °C. Use EASY-nLCTM 1200 nano to upgrade the UHPLC system, and the precolumn is a self-made precolumn Column (4.5 μm × seventy-five μm, 3 μm), the analytical column is a self-made analytical column (15 cm × one hundred and fifty μm, 1.9 μm). Use Q ExactiveTM series mass spectrometer for secondary mass spectrometry detection. The resolution of secondary mass spectrometry is set to 45000 (200 m/z), and the maximum capacity of C-trap is 5 × 10^4^, the maximum C-trap injection time is 86 ms, the peptide fragment fragmentation collision energy is set to 32%, and the threshold intensity is set to 1.2 × 10^5^, the dynamic exclusion range is set as 20 s, and the original data of mass spectrometry detection is generated.

### Proteome Data processing and bioinformatics Data Analysis

Use interproscan software to annotate GO and IPR functions (including Pfam, PRINTS, ProDom, SMART ProSite, PANTHER database), COG and KEGG carry out functional protein families and pathways for identified proteins Analysis (Jones P et al.,2014). Volcanic map analysis, cluster heat map analysis and pathway enrichment of GO, IPR and KEGG were conducted for DPE Analyze (Huang D W et al., 2009) and used STRING DB software to predict possible protein-protein interactions (http://STRING.embl.de/) (Franceschini et al., 2012).

### Single islets *β* cell preparation

The mice were anesthetized, and the abdominal cavity was opened. Collagenase V (10 mg/ml) was injected retrogradely from the pancreatic duct into pancreas. After digestion for 20 min at 37℃, the pancreatic tissue was removed and placed in pre-cooled Hank’s solution to terminate the digestion. The digested tissues were placed into petri dishes and structurally intact pancreatic islets were selected under a stereomicroscope. Selected pancreatic islets were then digested into single cell, washing, counting, and concentrating cells preparation for use according to10x Genomics Single Cell Protocols.

### Single-Cell RNA Sequencing (scRNA-seq)

The cell suspension was loaded into Chromium microfluidic chips with 3’ (v2 or v3, depends on project) chemistry and barcoded with a 10x Chromium Controller (10X Genomics). RNA from the barcoded cells was subsequently reverse-transcribed and sequencing libraries constructed with reagents from a Chromium Single Cell 3’ v2(v2 or v3, depends on project) reagent kit (10X Genomics) according to the manufacturer’s instructions. Sequencing was performed with Illumina (HiSeq 2000 or NovaSeq, depends on project) according to the manufacturer’s instructions (Illumina).

### Generation and Analysis of Single-Cell Transcriptomes

Fastp was used to perform basic statistics on the quality of the raw reads. Generally, celltanger count support FASTQ files from raw base call (BCL) files generated by Illumina sequencers as input file. For each gene and each cell barcode (filtered by CellRanger), unique molecule identifiers were counted to construct digital expression matrices. Secondary filtration by Seurat: A gene with expression in more than 3 cells was considered as expressed, and each cell was required to have at least 200 expressed genes. Then Secondary Analysis of Gene Expression, Global Analysis Between Samples, Enrichment analysis of cell/cell cluster markers was performed respectively.

### Directed differentiation of pancreatic islets

Undifferentiated human embryonic stem cell lines H1 and H9 and induced pluripotent cells UE005 were cultured using mTeSR1 (Stem Cells, Cat# 85850) in 6-well plates coated with Matrigel (Corning, Cat# 354277) in an incubator at 37 ℃ with 5% CO_2_. When the undifferentiated cells reached ∼70-80% confluence, treating with TrypLE Express (Thermo Fisher, Cat# 12605028) at 37 ℃ for 3-5 min. The released individual cells were washed with mTeSR1 and spun at 300 g rpm for 5 min. The resulting cell mass was resuspended in mTeSR1 medium supplemented with 10 μM Y-27632 (StemCell Technologies, Cat# 72307) and the single-cell suspension was seeded on Matrigel-coated 24-well plates at a ratio of ∼1.25 × 10^5^ cells/cm^2^. Cultures were fed daily with mTeSR1 medium, and differentiation was initiated 48 hours after seeding when the cells reached ∼90% fusion. Directed differentiation was carried out using the following 7-stage protocol, and then treated with 10 μM CDS or YOH for 7 days from S7D1. Phase 1 (S1, 3 days): Day 1 S1 medium + 100 ng/mL Activin A (StemCell Technologies, Cat# 78001) + 3 μM Chir99021 (Stemgent, Cat# 04-0004-10); Day 2 S1 medium + 100 ng/mL Activin A + 0.3 μM Chir99021; Day 3 S1 medium + 100 ng/mL Activin A

Phase 2 (S2, 2 days): S2 medium + 50 ng/mL FGF7 (StemCell, Cat# 78046)

Phase 3 (S3, 2 days): S3 medium + 50 ng/mL FGF7 + 1 μM retinoic acid (Sigma, Cat# R2625) + 0.25 μM SANT-1 (Sigma, Cat# S4572) + 100 nM LDN193189 (Reprocell, Cat# 40074) + 500 nM PdBU (Millipore, Cat# 524390) + 10 μM Y27632

Phase 4 (S4, 3 days): S4 medium + 50 ng/ml FGF7 + 20 ng/ml FGF2 (Med Chem Express, Cat# HY-P7004) + 0.1 μM retinoic acid + 0.25 μM SANT-1 + 100 nM LDN193189 + 100 nM PdBU + 10 μM Y27632 + 10 μM ALK5i II (Cell Guidance Systems, Cat# SM09-50)

Phase 5 (S5, 3 days): S5 medium + 50 ng/ml FGF7 + 20 ng/ml FGF2 + 0.05 μM retinoic acid + 0. 25 μM SANT-1 + 100 nM LDN193189 + 10 μM zinc sulfate (Sigma, Cat# Z0251) + 10 μM ALK5i II + 1 μM XXI (Millipore, Cat# 595790) + 20 ng/mL Betacellulin (Med Chem Express, Cat# HY-P7005) + 10 μg/ml Heparin (Sigma, Cat# H3149-500KU) + 1 μM T3 (Biosciences. Cat# 64245)

Phase 6 (S6, 5 days): S6 medium + 100 nM LDN193189 + 0.1 μM XXI + 10 μg/ml heparin + 2 μM R428 (Selleck Chem, Cat# S2841) + 10 μM Zinc Sulfate + 1 mM N-Acetylcysteine + 1 μM T3

Phase 7 (S7, 7-14 days): S7 medium + 1 mM N-acetyl cysteine + 10 μM zinc sulfate + 10 μM (±)-α-Tocopherol (Sigma, Cat# T3251) + 10 μg/ml heparin + 1 μM T3

### Analysis of gene expression

Total pancreatic islets RNA was isolated using the RNA extraction (Accurate Biology) accordance to the manufacturer’s instructions. The isolated RNA was reverse-transcribed with High-Capacity cDNA RT Kit and amplified with Takyon LowRox MasterMix dTTP Blue (Eurogentec). Quantitative PCR was performed by QuantStudio 1 Real-Time PCR System (Thermo Fisher Scientific, Cleveland, OH, USA). Relative expression of samples was adjusted for total RNA content and normalized to the mRNA expression levels of the β-actin or GAPDH. Data was analyzed using the 2-DDCt method (see key resources table for list of primers).

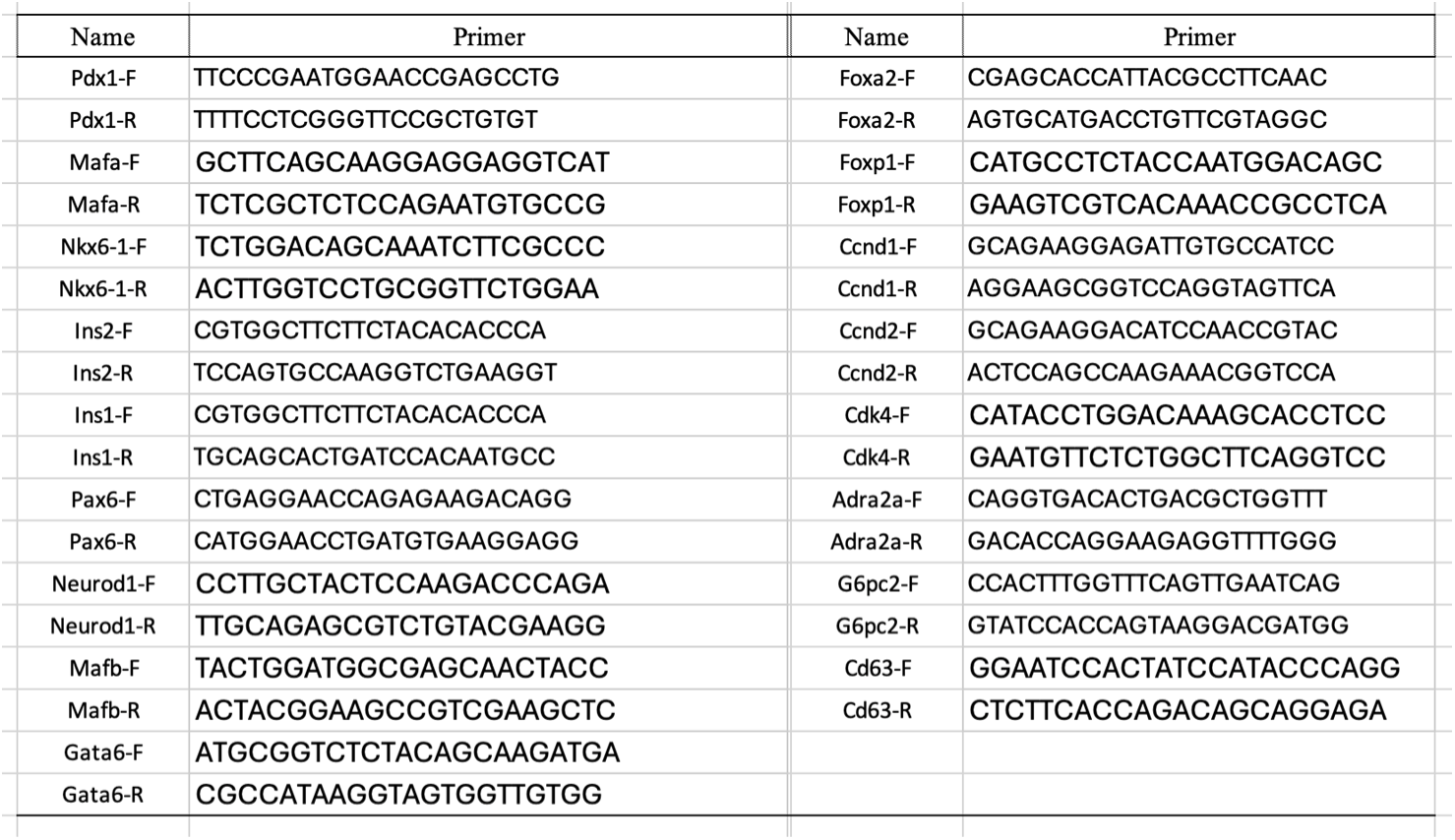

### General Procedure for development of the new compounds

All solvents and reagents were obtained from commercial suppliers and used without further purification. All non-aqueous reactions were run under an inert atmosphere (nitrogen or argon) with the rigid exclusion of moisture from reagents, and all reaction vessels were oven dried. All the reactions were monitored by thin-layer chromatography (TLC), carried out on silica gel plates (HSGF 254, Yantai Jiangyou Chemical, Yantai, China). Spots were visualized under UV at 254 nm. *Column chromatography* was carried out using silica gel (200–300 mesh, Qingdao Haiwan Specialty Chemicals Co. Ltd., Qingdao, China). ^1^H-NMR and ^13^C-NMR spectra were measured on a Bruker Avance Ⅲ 400 MHz NMR using deuterated chloroform (CDCl_3_) and deuterated dimethyl sulfoxide (DMSO-*d_6_*) as the solvent. Chemical shifts are expressed in δ (ppm). Abbreviations for peak patterns in NMR spectra: br = broad, s = singlet, d = doublet, t = triplet, q = quartet, dd=double doublet, m = multiplet. Coupling constants (J) are given in Hz. Low-resolution mass data were obtained with Agilent 1200 Quadrupole LC/MS. High-resolution mass spectroscopy (HRMS) was processed on a TOF instrument equipped with positive ionization mode of ESI. All HPLC data were obtained on the Agilent 1200 Series HPLC with autosampler, a quaternary pump, and UV-DAD detector. The column was Zorbax SB C18 column (4.6 mm × 150 mm, 5μm). Conditions were as follows: The mobile phase buffer was composed of 0.1% (v/v) TFA in H_2_O (buffer A) and ACN (buffer B). The total time of the detection method was 30 min. The separation gradient increases linearly from 10% buffer B to 90% buffer B during 0-30 min. The ratio of eluent was at 1.0 mL/min flow. Data were collected and analyzed with Agilent Chemstation software. The following abbreviations for solvents and reagents are used: tetrahydrofuran (THF), dichloromethane (DCM), *N,N*-dimethylformamide (DMF), ethyl acetate (EtOAc), (1-cyano-2-ethoxy-2-oxoethylidenaminooxy)dimethylamino-morpholino-carbenium hexafluorophosphate (COMU), bis(dibenzylideneacetone) palladium (Pd(dba)_2_), 1,1’-bis(diphenylphosphino)ferrocene (dppf).

Synthesis of compound **CDS479-2***^a^*

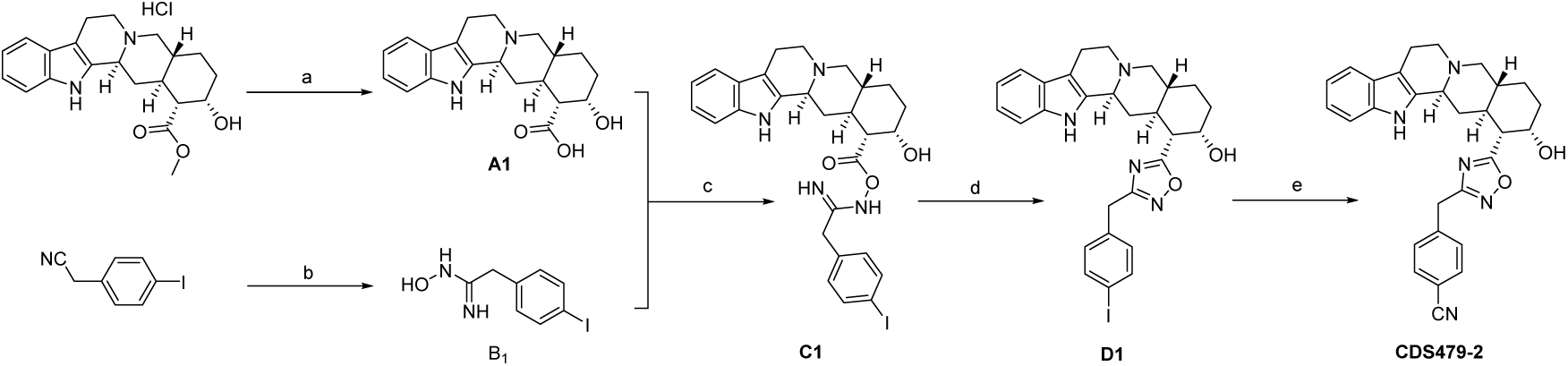

*^a^*Reagents and conditions: (a) LiOH·H_2_O, MeOH/THF/H_2_O, 0 °C∼r.t.; (b) Na_2_CO_3_, NH_2_OH·HCl, EtOH/H_2_O, r.t., 24h; (c) COMU, Et_3_N, DMF, r.t., 24h; (d) dioxane/toluene, 100 °C, 24h; (e) Pd(dba)_2_, dppf, Zn (CN)_2_, DMF, 80 °C, 6 h

#### (1R,2S,4aR,13bS,14aS)-2-hydroxy-1,2,3,4,4a,5,7,8,13,13b,14,14a-dodecahydroindolo [2’,3’:3,4] pyrido[1,2-b]isoquinoline-1-carboxylic acid (A1)

Lithium hydroxide monohydrate (5.04 g, 120 mmol) was added to a solution of yohimbine hydrochloride (7.81 g, 20 mmol) in MeOH/THF/H_2_O (90 mL/60 mL/30 mL) at 0 °C. After 30 min, the reaction was allowed to warm to room temperature and monitored by LC-MS until completion. The solvent was evaporated and the residue was dissolved in H_2_O (150 mL). Then the aqueous phase was acidified to pH = 6 with 5% HCl in an ice bath and let it sit overnight. The resulting precipitate was filtered and washed with water (100 mL) to afford **A1** as a white solid (5.9 g, yield 86%). ^1^H NMR (400 MHz, DMSO-*d*_6_) δ 10.81 (s, 1H), 7.33 (d, *J* = 7.7 Hz, 1H), 7.26 (d, *J* = 7.9 Hz, 1H), 7.03 – 6.96 (m, 1H), 6.95 – 6.88 (m, 1H), 4.13 (s, 1H), 3.29 (d, *J* = 11.1 Hz, 1H), 3.06 – 2.98 (m, 1H), 2.90 – 2.73 (m, 2H), 2.63 – 2.46 (m, 4H), 2.21 – 2.10 (m, 2H), 1.88 – 1.71 (m, 2H), 1.57 – 1.22 (m, 4H), 1.06 – 0.94 (m, 1H).

#### N-hydroxy-2-(4-iodophenyl) acetimidamide (B1)

Sodium carbonate (848 mg, 8 mmol) and hydroxylamine hydrochloride (695 mg, 10 mmol) were added to a solution of 2-(4-iodophenyl) acetonitrile (2.43 g, 10 mmol) in EtOH/H_2_O (20 mL/8 mL), then the mixture was stirred at room temperature for 24 hours. The solvent was evaporated, and the residue was dissolved in EtOAc (80 mL). The organic phase was washed with brine (20 mL × 2), dried by Na_2_SO_4_, filtered and concentrated. The crude product **B1** as a light yellow solid (2.77 g, yield >100%) was used directly for the next step without further purification.

#### N-(((1R,2S,4aR,13bS,14aS)-2-hydroxy-1,2,3,4,4a,5,7,8,13,13b,14,14a-dodecahydroindolo[2’,3’:3,4]pyrido[1,2-b]isoquinoline-1-carbonyl)oxy)-2-(4-iodophenyl)acetimidamide (C1)

Et_3_N (304 mg, 3 mmol) was added to a solution of **A1** (340 mg, 1 mmol) in DMF (5 mL) at room temperature. After 5 minutes, COMU (428 mg, 1 mmol) was added to the mixture and the mixture was stirred at room temperature for 20 minutes. Then crude product **B1** (552 mg, 2 mmol) was added to the mixture and the reaction was stirred at room temperature for 24 hours. The solvent was evaporated and the residue was dissolved in DCM/MeOH (100 mL/10 mL). The organic phase was washed with brine (20 mL × 3), dried by Na_2_SO_4_, filtered and concentrated. The crude residue was purified by column chromatography on silica gel (DCM/MeOH = 30:1) to give **C1** as a brown foam-like solid.

#### (1R,2S,4aR,13bS,14aS)-1-(3-(4-iodobenzyl)-1,2,4-oxadiazol-5-yl)-1,2,3,4,4a,5,7,8,13,13b,14,14a-dodecahydroindolo[2’,3’:3,4]pyrido[1,2-b]isoquinolin-2-ol (D1)

The intermediate **C1** was dissolved in anhydrous 1,4-dioxane/toluene (5 mL/10 mL) under nitrogen atmosphere, the mixture was heated at 100 ℃ for 24 hours. The solvent was evaporated and the residue was dissolved in DCM/MeOH (100 mL/10 mL). Then the organic phase was washed with brine (20 mL × 3), dried by Na_2_SO_4_, filtered and concentrated. The crude residue was purified by column chromatography on silica gel (DCM/MeOH = 30:1) to give **D1** as a brown foam-like solid (70 mg, total yield of two steps is 12%).

#### 4-((5-((1R,2S,4aR,13bS,14aS)-2-hydroxy-1,2,3,4,4a,5,7,8,13,13b,14,14a-dodecahydroindolo[2’,3’:3,4]pyrido[1,2-b]isoquinolin-1-yl)-1,2,4-oxadiazol-3-yl)methyl)benzonitrile (CDS479-2)

The compound **D1** (58 mg, 0.1 mmol) was dissolved in DMF (5 mL), then Pd(dba)_2_ (5.7 mg, 0.01 mmol), dppf (5.5 mg, 0.01 mmol) and Zn(CN)_2_ (11.7 mg, 0.1 mmol) were added under nitrogen atmosphere, the mixture was heated at 80 ℃ for 6 hours. The solvent was evaporated and the residue was dissolved in DCM/MeOH (100 mL/10 mL). Then the organic phase was washed with brine (20 mL × 3), dried by Na_2_SO_4_, filtered and concentrated. The crude residue was purified by column chromatography on silica gel (DCM/MeOH = 30:1) to give **CDS479-2** as a brown foam-like solid (33 mg, yield 70%). ^1^H NMR (400 MHz, DMSO-*d*_6_) δ 10.69 (s, 1H), 7.83 (d, *J* = 7.6 Hz, 2H), 7.60 (d, *J* = 7.6 Hz, 2H), 7.33 (d, *J* = 7.6 Hz, 1H), 7.24 (d, *J* = 8.0 Hz, 1H), 7.02 – 6.95 (m, 1H), 6.95 – 6.88 (m, 1H), 4.80 (d, *J* = 4.8 Hz, 1H), 4.28 (s, 2H), 3.99 (s, 1H), 3.29 – 3.22 (m, 1H), 3.07 (d, *J* = 12.0 Hz, 1H), 3.04 – 2.97 (m, 1H), 2.88 (d, *J* = 10.8 Hz, 1H), 2.81 – 2.70 (m, 1H), 2.64 – 2.51 (m, 2H), 2.22 – 2.06 (m, 3H), 1.82 – 1.73 (m, 1H), 1.72 – 1.62 (m, 1H), 1.57 – 1.44 (m, 2H), 1.39 – 1.30 (m, 1H), 1.09 – 0.98 (m, 1H). ^13^C NMR (100 MHz, CDCl_3_) δ 181.37, 168.21, 140.43, 136.11, 134.09, 132.77, 130.11, 127.46, 121.70, 119.65, 118.63, 118.28, 111.72, 110.92, 108.55, 67.19, 61.32, 59.91, 53.10, 46.15, 41.53, 38.35, 34.38, 32.64, 31.38, 23.38, 21.82. HPLC purity: >96%. HRMS(ESI): calcd for C29H30N5O2 [M + H]^+^, 480.2394; found, 480.2395.

### Calcium mobilization assay

Cells expressing α_2A_AR and Gα16 were seeded in 96-well plates at a density of 4×10^4^ cells/well and cultured overnight. The cells were loaded with 2 μmol/L Fluo-4 AM in Hanks’ balanced salt solution (HBSS, 5.4 mmol/L KCl, 0.3 mmol/L Na_2_HPO_4_, 0.4 mmol/L KH_2_PO_4_, 4.2 mmol/L NaHCO_3_, 1.3 mmol/L CaCl_2_, 0.5 mmol/L MgCl_2_, 0.6 mmol/L Mg_2_SO_4_, 137 mmol/L NaCl, 5.6 mmol/L D-Glucose, 250 μmol/L Sulfinpyrazone, pH 7.4) at 37 ℃ for 45 min. After a thorough washing, 50 μL assay buffer containing compound was added. After incubation at 37°C for 10 min, 25 μL assay buffer containing agonists was dispensed into the well, using a FlexStation III microplate reader (Molecular Devices, Sunnyvale, CA, USA) and intracellular calcium change was recorded with an excitation wavelength of 485 nm and emission wavelength of 525 nm. IC_50_ values for each curve were calculated by Prism 5.0 software (GraphPad Software).

### Molecular Docking Study

The Crystal structure of compound RSC-bound alpha2A adrenergic receptor complex (PDB ID: 6KUX) and Yohimbine was chosen for molecular docking study. SchrodingerSuite (Windows v. 2018-1) was used as the docking software. The receptor was prepared by the Protein Preparation Wizard, and the ligand was processed by LigPrep module. The docking results were analyzed by the Ligand Docking module. The 3D interaction graphic was created by PyMOL software.

### Constructs of alpha2A-Fab-Glue complex

To stabilize the inactive alpha2A structure and increase its molecular weight, the alpha2A-Fab-glue protein fusion construct was employed. First, a fusion protein, mBRIL, was introduced into alpha2A to replace ICL3, providing structural rigidity and minimizing interference with the receptor, while simultaneously binding with Fab due to its anti-mBRIL characteristic. Another component, Glue, consists of NbFab-linker1-E3-linker2-NbALFA, with NbFab binding to Fab, and E3 and NbALFA binding to K3 and ALFA, respectively, which were extended after H8. Consequently, this forms a stable heterotrimer construct, enabling the inactive state of alpha2A with antagonist to be successfully captured with cryo-EM.

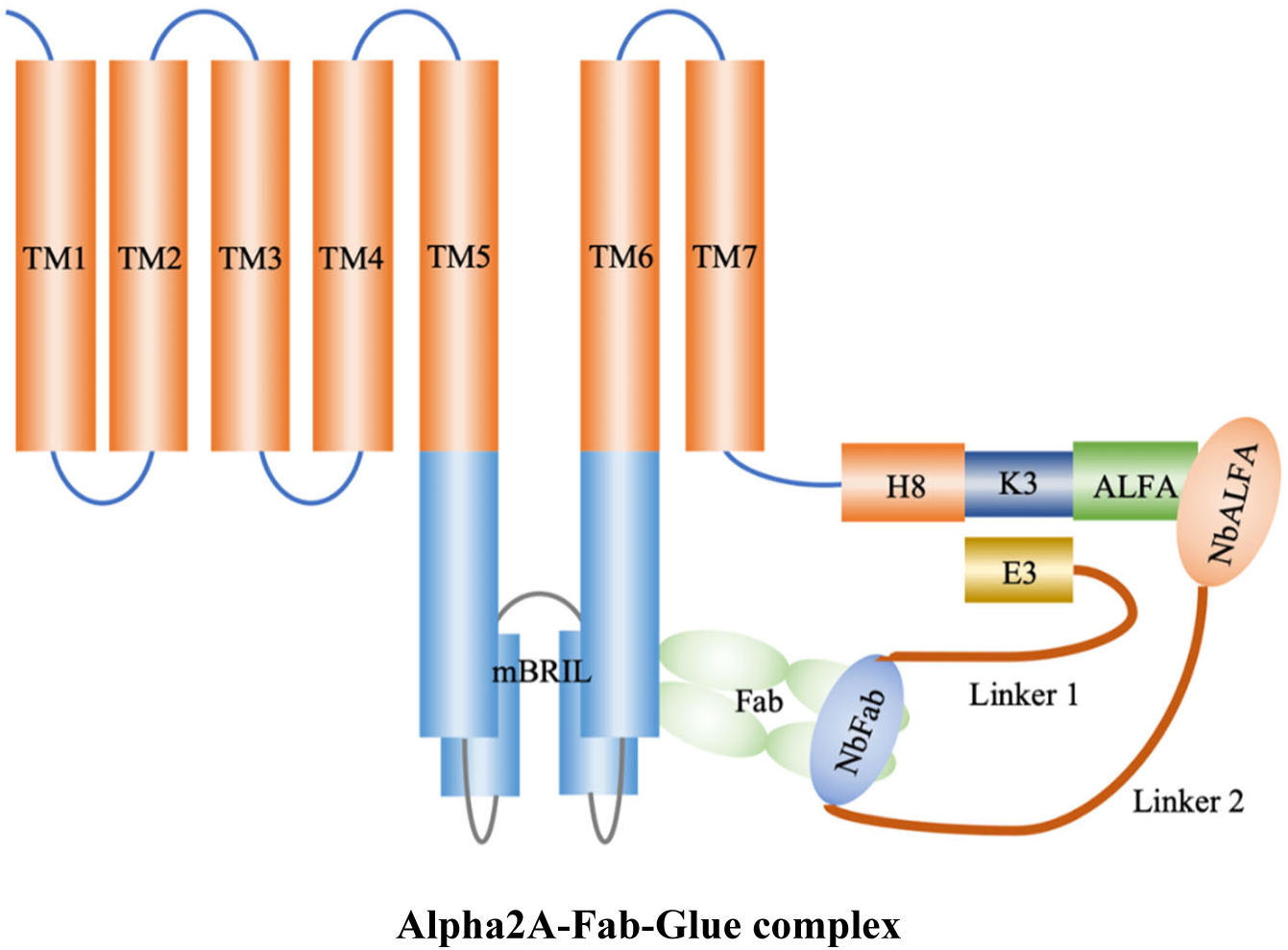

### Expression, formation, and purification of alpha2A-Fab-Glue complex

The coding sequence of alpha2A-mBRIL-K3-ALFA was cloned into pFastBac vector with a hemagglutinin (HA)signal and a Flag peptide tag at N-terminal, and a C-terminal 8×His tag. The plasmid was transfected into DH10Bac to produce recombinant baculovirus with the Bac-to-Bac system (Invitrogen). The Sf9 cells were infected by the baculovirus at a cell density of 4 × 10^6^ cells ml^-1^, and then cultured at 27°C, shaking at 110 rpm, for 48h. After cells were lysed with lysis buffer (10mM HEPEs PH7.5, 1mM PMSF) at 4 °C for 30min, the membrane was homogenized in solubilization buffer (20 mM HEPES pH 7.5, 500 mM NaCl, 10% (w/v) glycerol, 1% (w/v) n-dodecyl-B-D-maltoside (DDM, Anatrace Cat# D310) and 20µM CDS at 4 °C for 2h. The alpha2A-mBRIL-K3-ALFA was purified by Ni-NTA chromatography, washed with buffer containing 20mM HEPEs, 150mM NaCl, 0.03% (w/v) DDM, 0.01% (w/v) Lauryl maltose neopentyl glycol glycol (LMNG, Anatrace Cat# NG310), 0.001% (w/v) cholesteryl hemisuccinate (CHS, Sigma Cat# C6512), 20µM CDS, and 20mM imidazole, ended up with elution containing 20mM HEPEs, 150mM NaCl, 0.03% (w/v) DDM, 0.01% (w/v) LMNG, 0.001% (w/v) CHS, 20µM CDS, and 300mM imidazole.

The coding sequences of Glue and Fab were cloned into the pET-22b (+) vector with an N-terminal pelB signal peptide and a C-terminal 6×His tag. After plasmids were translated into E. coli BL21(DE3), the cells were cultured in LB medium with 1‰ (w/v) ampicillin at 37 °C, shaking at 220 rpm to OD_600_ = 0.8, and then induced with 1mM IPTG at 16°C, shaking at 150 rpm, for 24h. Cells were disrupted by sonication in NH buffer which contains 20 mM HEPES (pH 7.5) and 150 mM NaCl. The Fab and glue were purified by Ni-NTA chromatography, washed with NH buffer with 20mM imidazole, ended up with 300mM imidazole NH buffer elution.

After concentrated to 2 mg ml^-1^ with 10-kDa ultra centrifugal filter (Millipore), the Fab and glue were mixed with alpha2A-mBRIL-K3-ALFA for heterotrimer complex formation, at 4 °C for 2h, also with 2mM CaCl_2_ and M1 resin. The next step was detergent exchanging process with different proportions of buffer A (20mM HEPEs, 150mM NaCl, 0.03% (w/v) DDM, 0.01% (w/v) LMNG, 0.001% (w/v) CHS, 20µM CDS, and 2mM CaCl_2_) and buffer B (20mM HEPEs, 150mM NaCl, 0.1% (w/v) LMNG, 0.01% (w/v) CHS, 20µM CDS, and 2mM CaCl_2_) until the DDM was taken placed by LMNG. And then, the complex was eluted with buffer containing 20mM HEPEs, 150mM NaCl, 0.001% (w/v) LMNG, 0.0001% (w/v) CHS, 20µM CDS, 5mM EDTA and 200 µg ml^-1^ Flag peptide. After concentrated to 500µl with 50-kDa ultra centrifugal filter (Millipore), the complex was further purified with AKTA Superdex 200 Increase 10/300 GL column (Cytiva) preequilibrated with buffer containing 20mM HEPEs, 150mM NaCl, 0.001% (w/v) LMNG, 0.0001% (w/v) CHS, 20µM CDS to obtain a relatively pure trimer. Finally, the fractions were concentrated to 5mg ml^-1^ with 50-kDa ultra centrifugal filter for cryo-EM.

### Cryo-EM sample preparation and data collection

The gold film (UltraAuFoil/QuantiFoil, 300 mesh, R1.2/1.3, Quantifoil Micro Tools GmbH, DE) or amorphous alloy film (300 mesh, R1.2/1.3, Zhenjiang Lehua Electronic Technology Co., Ltd.) was glow discharged with air for 40 s at 15 mA at easiGlow™ Glow Discharge Cleaning System (PELCO, USA) or Tergeo Plasma Cleaner (PIE SCIENTIFIC, USA). 3 μL purified complex sample was dropped onto the grid and then blotted for 3-4 s with blotting force 0 and plunged into liquid ethane cooled by liquid nitrogen using Vitrobot Mark IV (Thermo Fisher Scientific, USA). Cryo-EM datasets were collected with the 300 kV Titan Krios Gi3 microscope. The raw movies were collected by Gatan K3 BioQuantum Camera at the magnification of 105000, with a pixel size of 0.85 Å. Inelastically scattered electrons were excluded by a GIF Quantum energy filter (Gatan, USA) using a slit width of 20 eV. The movies were acquired with the defocus range of −1.2 to −2.0 μm with a total exposure time of 2.5 s fragmented into 50 frames and with a dose rate from 15.34∼17.55 e/pixel/s. SerialEM was used for semi-automatic data acquisition.

### Data processing, model building and refinement

The image stacks were collected and subjected to motion correction using MotionCor2. Contrast transfer function parameters were estimated by CTFFIND4, implemented in RELION4.0 or Patch CTF in cryoSPARC. Particles were auto-picked from micrographs by RELION/cryoSPARC and then subjected to several rounds of 2D classification using cryoSPARC. Selected particles with an appropriate 2D average from 2D classification were further subjected to cryoSPARC to generate *Ab-initial* model. In the next step, 2 rounds of 3D classification were conducted in RELION. Eventually, particles with high-resolution 3D average were selected from 3D classification, which resulted in a map with an initial resolution of near-atomic level by

RELION. The refined particles were subjected to CTF refinement to update per-particle defocus and per-micrograph astigmatism. Furthermore, the particles are polished by RELION. Polished particles are imported into cryoSPARC to performed NU-refinement, a final map with resolution 3.34 Å were generated after postprocessing determined by gold-standard Fourier shell correlation using the 0.143 criteria. The final map was sharped and masked by deepEMhancer. The local resolution map was calculated from cryoSPARC using two unfiltered half maps. Released A2AR structure (PDB: 6kux) was using as a template for model building. Model docking was carried out using Chimera. Manually adjustment and rebuilding were performed with COOT. The model was repeatedly refined using real space refinement module in Phenix. The software UCSF ChimeraX was used to prepare the molecular graphics figures. The final model statistical analysis was performed by Phenix.

